# Global proteomics of *Ubqln*2-based murine models of ALS

**DOI:** 10.1101/2020.02.22.956524

**Authors:** Alexandra M. Whiteley, Miguel A. Prado, Stefanie A.H. de Poot, Joao A. Paulo, Marissa Ashton, Sara Dominguez, Martin Weber, Hai Ngu, John Szpyt, Mark P. Jedrychowski, Amy Easton, Steven P. Gygi, Thimo Kurz, Mervyn J. Monteiro, Eric J. Brown, Daniel Finley

**Author notes:** These authors contributed equally (Alexandra M. Whiteley, Miguel A. Prado).

## Abstract

Familial forms of neurodegenerative diseases commonly involve mutation of aggregation-prone proteins or components of the protein degradation machinery that act on aberrant proteins. *Ubqln2* encodes a member of the UBL/UBA family of proteasome shuttle factors that is thought to facilitate proteasomal degradation of substrates, and mutation of this gene results in a familial form of ALS/FTD in humans. How *Ubqln2* dysfunction leads to neurodegeneration, however, remains uncertain. We undertook a comprehensive study to identify proteomic changes upon *Ubqln2* perturbation in multiple murine models of *Ubqln2*-mediated neurodegenerative disease. By performing quantitative multiplexed proteomics on neural tissues of affected animals, we identified a small group of proteins whose abundance is tightly linked to UBQLN2 function: the ubiquitin ligase TRIM32 and two retroelement-derived proteins, PEG10 and CXX1B. Further studies using cultured cells of human origin, including induced neurons, found similar changes in protein abundance upon *Ubqln2* loss, and pulse-chase studies suggested that PEG10 and TRIM32 are direct clients of UBQLN2. In conclusion, our study provides a deep understanding of the proteomic landscape of ALS-related *Ubqln2* mutants and identifies candidate client proteins that are altered *in vivo* in disease models and whose degradation is promoted by UBQLN2.

## Introduction

Many progressive neurodegenerative diseases feature an accumulation of protein aggregates that are thought to promote or cause disease through the progressive loss of neuronal function (Ross and Poirier, 2004; Wood et al., 2003). Familial genetic variants often lead to conformational changes that cause proteins to misfold, thereby contributing to the development of disease, such as in the case of TDP-43-driven familial Amyotrophic Lateral Sclerosis (fALS; Sreedharan et al., 2008). In other cases, mutations in components of the protein degradation machinery itself such as Optineurin and P62/SQSTM1 result in aggregate formation and disease (Fecto et al., 2011; Maruyama et al., 2010). Amyotrophic lateral sclerosis (ALS), a progressive and fatal motor neuron disease that typically presents in the fourth to seventh decades of life and affects upwards of 30,000 adults in the United States, is characterized by the presence of protein inclusions in affected motor neurons (Blokhuis et al., 2013). fALS can similarly be caused by mutation of aggregation-prone proteins, such as TDP-43, SOD1, and C9ORF72, or of genes involved in protein degradation, such as VCP, Optineurin, and Ubiquilin2 (UBQLN2; Kiernan et al., 2011).

UBQLN2 belongs to a large family of proteins that link proteasomes with ubiquitinated proteins through a ubiquitin-like domain (UBL) that binds to the proteasome, and a ubiquitin-binding domain (UBA) that binds to a variety of mono- or polyubiquitinated proteins (Chen et al., 2019; Kleijnen et al., 2003; 2000; Zhang et al., 2008). As such, Ubqlns and other UBL/UBA proteins facilitate proteasomal degradation of ubiquitinated proteins. Our understanding of Ubqln biology is largely based on studies of the yeast homolog Dsk2 (Biggins et al., 1996; Elsasser and Finley, 2005; Liu et al., 2009; Medicherla et al., 2004), the UBL domain of which is recognized by subunits Rpn10, Rpn13, and Rpn1 of the proteasome (Elsasser et al., 2002; Finley, 2009; Walters et al., 2002; Shi et al., 2016). Mammalian Ubqlns appear to bind the proteasome via the orthologous subunits (Chen et al., 2016; 2019; Walters et al., 2002). The UBA domain of Dsk2 and mammalian Ubqlns has affinity for ubiquitin chains (Rao and Sastry, 2002; Saeki et al., 2002; Wilkinson et al., 2001) but little or no preference for particular ubiquitin linkages (Funakoshi et al., 2002; Harman and Monteiro, 2019; Hjerpe et al., 2009; Seok Ko et al., 2004; Sims et al., 2009; Zhang et al., 2008). However, for some clients, Ubqlns may bind their non-ubiquitinated forms and instead promote ubiquitination through recruitment of a ligase (Dao et al., 2018; Itakura et al., 2016).

While Dsk2 has been implicated in the large-scale proteasomal degradation of ubiquitinated protein (Funakoshi et al., 2002; Saeki et al., 2002; Tsuchiya et al., 2017), Ubqlns have been implicated in the degradation of specific classes of proteins, most strikingly hydrophobic and/or transmembrane proteins (Suzuki and Kawahara, 2016), aggregates (Hjerpe et al., 2016), and mislocalized mitochondrial proteins (Itakura et al., 2016; Whiteley et al., 2017). While these reports implicate different potential functions of Ubqlns, some of their apparent functional diversity may be due to the multiplicity of Ubqln isoforms. There are 6 *UBQLN* genes expressed in humans: *UBQLN1*, which is expressed throughout the body; *UBQLN2*, which is enriched in neuronal tissues and muscle; *UBQLN 3*, *5*, and *L* which are expressed only in testes; and *UBQLN4*, which is induced in many cells upon nutrient limitation (Marín, 2014). While all *UBQLN* genes have a similar domain organization featuring UBL and UBA domains, many also contain unique motifs that could potentially confer client specificity or functional specialization. For example, UBQLN4 possesses a non-canonical LC3-binding motif (Lee et al., 2013) which is thought to confer a unique ability of the Ubqln family to promote autophagic protein degradation (N’Diaye et al., 2009a; 2009b; Rothenberg et al., 2010; Şentürk et al., 2019). In addition, UBQLN2 contains a proline-rich repeat (the ‘PXX-domain’), which may regulate client specificity (Gilpin et al., 2015), proteasome binding (Chang and Monteiro, 2015), and liquid-liquid phase separation into stress granules (Alexander et al., 2018; Dao et al., 2018; Sharkey et al., 2018).

Mutations in *UBQLN2* cause a heritable, X-linked, dominant form of ALS with frontotemporal dementia (fALS/FTD) in humans (Deng et al., 2011). This discovery has been corroborated through the use of mutant transgenic mouse models that mimic *in vivo* symptoms of motor neuron disease and exhibit protein aggregates in the brain (Chang and Monteiro, 2015; Hjerpe et al., 2016). Although the initially identified mutations were all located in the PXX domain of *UBQLN2* (Deng et al., 2011), recent studies have also identified patients with mutations adjacent to (Teyssou et al., 2017; Williams et al., 2012) and completely outside of (Daoud et al., 2012; Synofzik et al., 2012) this region. The molecular mechanism of neurodegenerative disease due to *UBQLN2* mutation remains unknown but could involve progressive accumulation of as-yet-unidentified Ubqln target proteins. However, there have been no published comprehensive proteomic studies in any relevant model systems to identify target proteins in an unbiased fashion.

Ubqlns have been shown to regulate the abundance of individual proteins linked to neurodegenerative disease, including Huntingtin (Hjerpe et al., 2016; Rutherford et al., 2013; Wang et al., 2006), Aß (Ayadi et al., 2012), and presenilins (Mah et al., 2000). Furthermore, UBQLN2 colocalizes with TDP-43 inclusions of familial ALS not only when UBQLN2 is mutated (Deng et al., 2011), but also in familial ALS characterized by an ALS-associated FUS mutation (Williams et al., 2012) or mutant TDP-43 overexpression (Deng et al., 2011). Like other Ubqlns, UBQLN2 oligomerizes (Ford and Monteiro, 2006) and is capable of phase separation which could potentially contribute to disease by influencing protein degradation pathways and even nucleating protein aggregation within the cell (Alexander et al., 2018; Dao et al., 2018; Sharkey et al., 2018). Thus, perturbation of UBQLN2 function may promote neurodegeneration by compromising the proteasomal degradation of proteins known to cause ALS, such as TDP-43. Alternatively, *Ubqln2* mutations may lead to neurodegeneration through the dysregulation of unrelated, unknown proteins, through a general attenuation of protein quality control, or through an inherent toxicity that is unrelated to the normal function of UBQLN2 in protein degradation.

In this study, we have performed a comprehensive proteomic analysis of multiple animal models of *Ubqln2*-driven neurodegenerative disease. In doing so, we have identified proteins whose abundance changes upon *Ubqln2* perturbation and thus are not only linked to Ubqln function but may also be involved in the development of neurodegenerative disease. Furthermore, through the utilization of pathway analysis we were able to identify groups of proteins that were also functionally linked to *Ubqln2* dysfunction. By comparing results from multiple *Ubqln2*-dependent models of ALS, we were able to focus our results on a select group of proteins which were commonly altered upon different forms of *Ubqln2* perturbation and assess their relationship to UBQLN2. From this global search we have identified exceptionally responsive UBQLN2 clients in the brain, notably the E3 ubiquitin ligase TRIM32 and the retroelement product PEG10. Unexpectedly, we also found UBQLN2 to strongly protect from degradation the Gag-like protein CXX1B, which is expressed from the same family of retroelement genes as PEG10–the Mar family. In conclusion, our results elucidate the proteomic changes that occur throughout the disease course of Ubqln-mediated neurodegeneration in murine model systems, revealing potential pathways of disease development as well as novel putative clients of UBQLN2.

## Results

### *Ubqln2^-/-^* animals develop age-dependent neuromotor defects

*Ubqln2*-deficient mice were generated by flanking the one exon of murine *Ubqln2* with *loxP* sites (Figure 1A) and expressing Cre-bearing plasmids in flox-bearing C57BL/6 ES cells. Complete loss of UBQLN2 protein was confirmed by Western blot of spinal cord tissue from WT and *Ubqln2^-/-^* animals (Figure 1B), whereas UBQLN1 protein levels were unaffected by the loss of UBQLN2 (Figure 1B). At weaning, there were no gross motor defects in *Ubqln2^-/-^* pups compared to their WT littermates. Therefore, to examine how these animals are affected by *Ubqln2* loss, 10-14 month old male WT and *Ubqln2^-/-^* animals (Supplemental Table 2) were tested in a battery of neuromotor exams. *Ubqln2^-/-^* animals consistently displayed more center beam breaks in open field tests (Figure 1C), indicating hyperactivity or a lack of fear/anxiety during repeated testing. *Ubqln2^-/-^* mice also exhibited increased hind limb clasping (Figure 1D), decreased wire hang time (Figure 1E) and increased footslips on both large and medium balance beams (Figure 1F) as compared to WT littermates despite no differences in weight (Supplemental Table 2). *Ubqln2^-/-^* mutants demonstrated transient changes in active avoidance of aversive stimuli (Supplemental Figure 1A-B) and mild defects in fear conditioning (Supplemental Figure 1C-E). Despite these phenotypes, there were no detectable changes in the number of neurons, markers of neuroinflammation, or ubiquitin-containing aggregates in *Ubqln2^-/-^* spinal cord (Supplemental Figure 2). From these results, we concluded that these animals have an intermediate, age-dependent neuromotor disease more severe than that seen in *Ubqln2* knock-in mice containing a mutant murine *Ubqln2* allele corresponding to the disease-causing P506T mutation (Hjerpe et al., 2016) and less severe than that of mice overexpressing a disease-causing mutant human P497S *UBQLN2* allele (Le et al., 2016).

**Figure 1:**
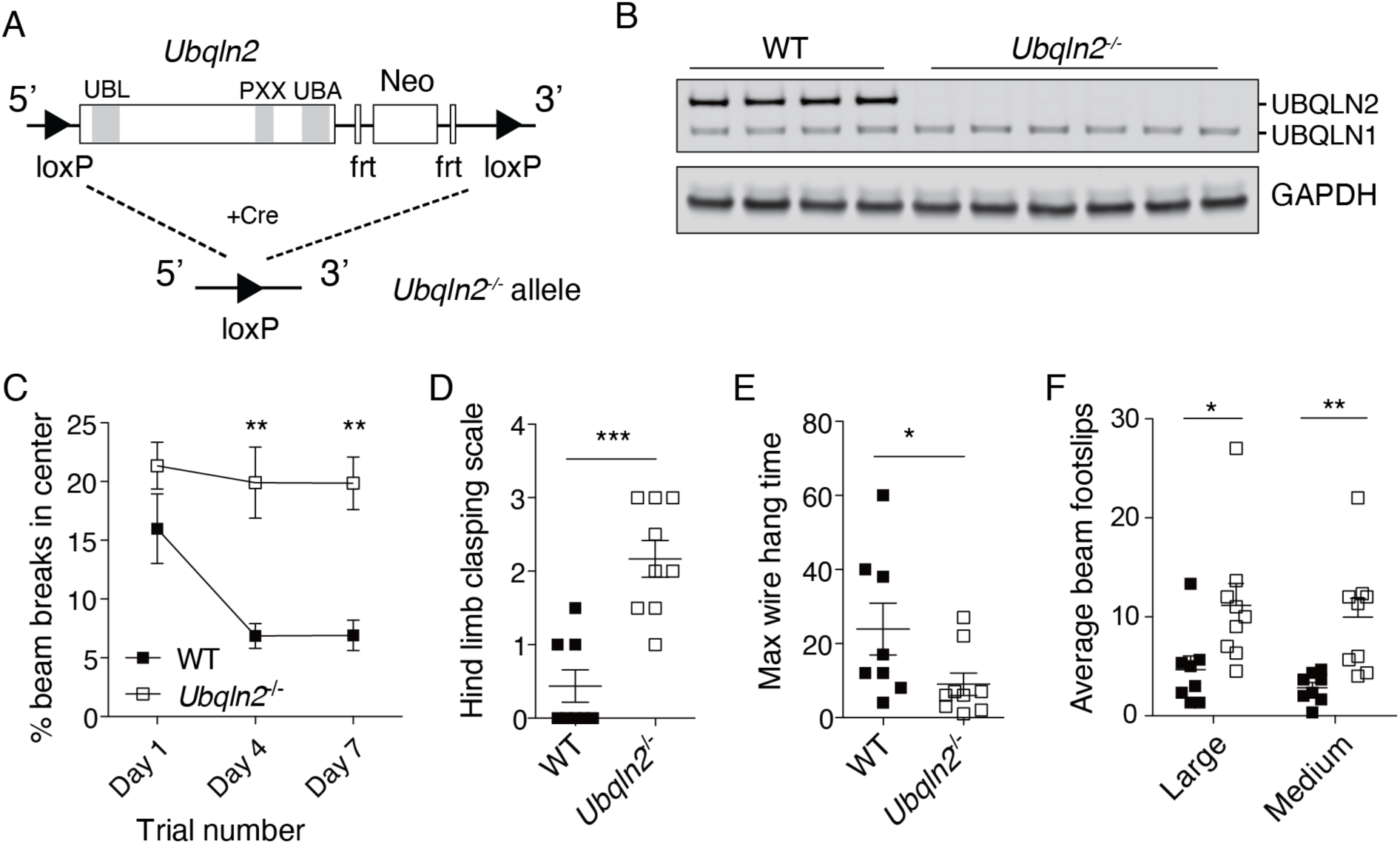
Neurodegenerative phenotype of *Ubqln2^-/-^* mice. (A) Graphical representation of genetic *Ubqln2* deletion. Functional domains are shown in grey and annotated above graphics. Neo=neomycin resistance cassette. Frt=Flp recombination sites. (B) Western blot demonstrating loss of UBQLN2 protein, but not UBQLN1 protein, in spinal cord samples of 5-6-month old mice. (C-F) Age-matched 10-14-month old male mice were tested in a battery of neuromotor exams. (C) *Ubqln2^-/-^* mice were more active over multiple sessions in an open field test as measured by % center beam breaks during the course of each session. X-axis shows the repeated locomotor sessions separated by three days. Statistical significance was determined with repeated measures ANOVA. (D) *Ubqln2^-/-^* mice exhibit increased hind limb clasping upon tail suspension and (E) decreased wire hang time compared to littermates. (F) *Ubqln2^-/-^* mice have increased footslips when on large and medium balance beam. N=8-9 mice per genotype. Statistical significance for (D-F) was determined with unpaired, two-tailed Student’s T test.

### *Ubqln2^-/-^* tissues demonstrate proteomic changes prior to development of neuromotor defects

Our main interest in *Ubqln2^-/-^* animals was to examine neural tissue for possible changes to the proteome. Therefore, five unique brain hemispheres of each genotype were isolated from mice at 12-16 months of age for proteomic analysis (Supplemental Table 2). Homogenized protein samples were digested and labelled with tandem mass tags (TMT) for multiplexed quantitative proteomic analysis using LC-SPS-MS3 analysis. Both hierarchical clustering (Figure 2A) and principal component analysis (Figure 2B) of proteomics data demonstrated distinct separation of samples based on *Ubqln2* expression. Changes to individual proteins are highlighted in Figure 2C; there was a substantial upregulation of the E3 ubiquitin ligases TRIM32 and RNF112 (also known as neurolastin or ZNF179), both of which are known to regulate neuronal development and function (Lomash et al., 2015; Sato et al., 2011). Mutation of TRIM32 NHL domains causes the neuromuscular disease Limb-Girdle muscular dystrophy type 2H (LGMD2H), whereas mutation in the BBOX domain of TRIM32 results in the ciliopathic disorder Bardet-Biedl Syndrome (Chiang et al., 2006; Kudryashova et al., 2005). TRIM32 ubiquitinates dysbindin (Locke et al., 2009) and actin (Cohen et al., 2012), while the substrates of RNF112 are unknown. Finally, there was a minor but highly significant increase in the mitochondrial protein ATAD1 (Figure 2C). ATAD1 and its yeast homolog Msp1 remove mislocalized tail-anchored proteins from the mitochondrial membrane (Chen et al., 2014; Okreglak and Walter, 2014); ATAD1 upregulation has also been observed upon loss of *Ubqln1* (Whiteley et al., 2017), which facilitates degradation of mislocalized mitochondrial proteins (Itakura et al., 2016). Thus, the upregulation of ATAD1 in *Ubqln2*-deficient tissues implicates a shared pathway linking UBQLN1, UBQLN2, mislocalized mitochondrial proteins, and ATAD1 protein abundance.

**Figure 2:**
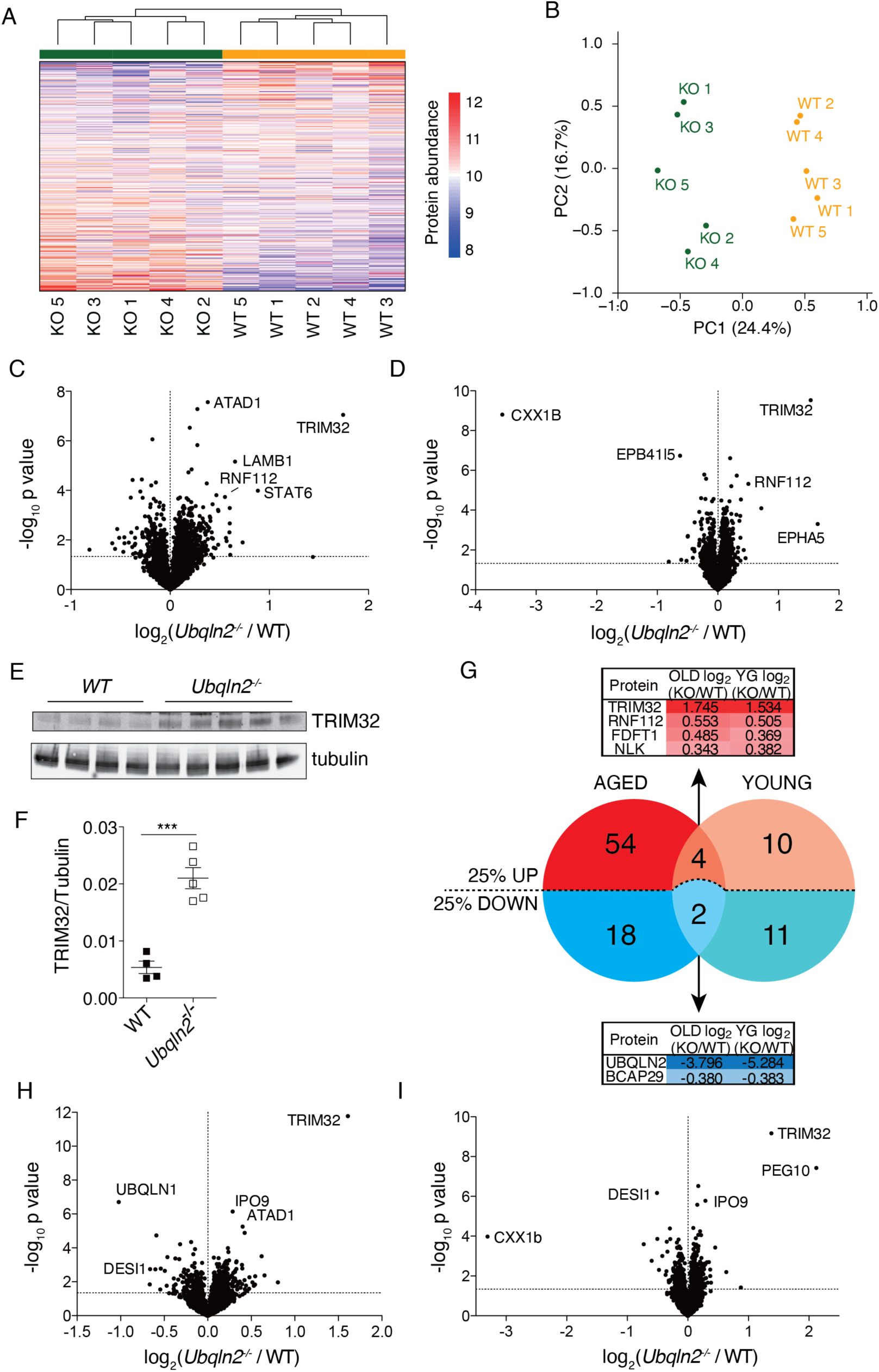
Proteomics of brain and spinal cord reveals proteins changed upon *Ubqln2* loss. (A-C) Brain hemispheres from 12-16-month old WT (N=5) and *Ubqln2^-/-^* (‘KO’, N=5) mice were isolated and analyzed by TMT. (A) Heatmap of brain hemisphere samples showing hierarchical clustering using a Euclidean distance of WT (yellow) and KO (green) samples. (B) Principal component analysis (PCA) of brain samples. WT are shown in yellow and KO in green. (C) Volcano plots with log2 ratios of mean protein abundances of all quantified proteins, with increasing significance on the y-axis (p value of 0.05 noted with y-axis dashed line). (D) Brain hemispheres were dissected from younger mice (5-6 months old) for TMT analysis. N=6 *Ubqln2^-/-^* and N=4 WT mice in both experiments. (E) Spinal cord of 5-6 month old animals was lysed in urea buffer and prepared for Western blot of TRIM32 and tubulin. (F) TRIM32 quantification normalized to tubulin. Statistical significance was determined with two-tailed, unpaired Student’s T test. (G) Venn diagram of proteins from aged (C) and young (D) brain hemispheres that are up- or down-regulated more than 25 % (p value < 0.05). Four proteins were increased at least 25% in both aged and young tissues (top, red) and two proteins were decreased at least 25% in both (bottom, blue). Tables highlight the log2 ratios of each named protein from Supplemental Table 3. (H) Hippocampus (N=5 samples from three biological replicates, for both genotypes) was isolated from 5-6 month old mice and (I) lumbar spinal cord (N=6 *Ubqln2^-/-^* and 4 WT) were isolated from 4-month old animals for TMT analysis. Information on total number of proteins identified and quantified in each TMT from this study is included in Supplementary Tables 3 and 5.

Because samples were taken for proteomics from aged animals that exhibited *in vivo* neuromotor defects, changes to the proteome could reflect chronic disease as opposed to an acute disturbance due to *Ubqln2* loss. We were particularly interested in observing the early proteomic changes that predate overt phenotypic effects of *Ubqln2* loss. Therefore, proteomics was also performed on brain hemispheres of animals at 5-6 months of age, when the *in vivo* effects of disease had not reached that of aged animals and did not yet achieve statistical significance (Supplemental Figure 3). As in aged brains, a small number of proteins were dramatically changed in *Ubqln2^-/-^* hemispheres of young mice, many of which were shared with older brain samples. As in aged animals, the E3 ubiquitin ligases TRIM32 and RNF112 were upregulated, particularly TRIM32 (Figure 2D), which was also confirmed by western blotting (Figure 2E-F). While most affected proteins were upregulated in *Ubqln2^-/-^* tissues compared to their WT counterparts, there was a dramatic loss in *Ubqln2^-/-^* brains of the protein CXX1B, which was likewise confirmed by Western blotting (Supplemental Figure 4A). CXX1B, also known as FAM127 or RTL8, is a Gag-like protein (Supplemental Figure 4) of unknown function expressed from specific members of the Ty3/Gypsy family of endogenous LTR-derived retroelements (Brandt et al., 2005).

Proteomic hits from brain hemisphere samples could be divided into three groups: those that were similarly affected by *Ubqln2* loss regardless of age, as expected of UBQLN2 client proteins; those showing more dramatic changes in aged *Ubqln2^-/-^* animals; and those showing alteration in young but not aged *Ubqln2^-/-^* animals (Figure 2G). The E3 ligases TRIM32 and RNF112, as well as the metabolic regulator FDFT1 and Nemo-like kinase (NLK) did not significantly change upon aging of *Ubqln2^-/-^* animals, indicating that these are early and persistent changes to the proteome upon *Ubqln2* loss. In addition, the B cell receptor-associated protein 29 (BCAP29) was downregulated in both aged and young brains (Figure 2G). LAMB1 and ATAD1 were only significantly upregulated in aged tissues and appear to represent sensitive indicators of the progression of neuromotor disease in *Ubqln2^-/-^* animals. EPHA5 and EPB41L5, on the other hand, were both altered in young tissues and unaffected by *Ubqln2* loss in aged animals.

To further define the effects of Ubqln2 on tissue proteomes relevant to the development of ALS, we isolated particular regions of CNS tissue known to express high levels of UBQLN2 and to play a role in disease. Hippocampus and lumbar spinal cord were isolated from *Ubqln2^-/-^* mice between 4 and 6 months of age (Supplemental Table 2), before they exhibited significant motor defects (Supplemental Figure 3), and tissue was prepared for proteomics (Figure 2H-I). Most notably, TRIM32 was found to be elevated in these tissues as a result of the *Ubqln2* null mutation, similarly to the brain hemisphere samples. ATAD1 was elevated upon *Ubqln2* loss in the hippocampus as had been observed in brain hemisphere samples. Unlike spinal cord and brain, hippocampus showed a clear decrease in UBQLN1 protein expression (Figure 2H), which may reflect a difference among tissues in their response to *Ubqln2* loss. In the case of spinal cord, there was also a dramatic loss of CXX1B, comparable to that described above for young brain. In addition, spinal cord revealed a strong upregulation of the protein PEG10, which, like CXX1B, is expressed from specific members of the Ty3/Gypsy family of retroelements (Brandt et al., 2005).

PEG10 is expressed primarily in two forms, PEG10-RF1 and PEG10-RF1/2, which correspond to the retroviral gene products Gag and Gag-Pol, respectively (Supplemental Figure 4B-C). In retroviruses, the Pol element encodes a retroviral protease, reverse transcriptase, and integrase, whereas PEG10-RF1/2 lacks reverse transcriptase activity due to deletions within this element. PEG10-RF1/2 is, like the ancestral Gag-Pol protein, generated by a highly efficient translational frameshift between the Gag-like and Pol-like elements (Clark et al., 2007; Shigemoto et al., 2001). While in retroviruses the Pol element is subsequently cleaved from the Gag element, this proteolytic cleavage is at best weak in PEG10 (Abed et al., 2019). Instead, the “Gag-only” form PEG10-RF1 is produced when the frameshift element is bypassed and the proximal stop codon is engaged (Supplemental Figure 4C).

PEG10 upregulation promotes proliferation and invasiveness of cancer cells (Akamatsu et al., 2015), and deletion in mouse models causes lethality due to placentation defects (Ono et al., 2006). Recently, it was found that PEG10 regulates mRNA abundance in murine trophoblast cells, thereby promoting their differentiation in culture (Abed et al., 2019). Furthermore, purification of PEG10-RF1 yielded virus-like particles (Abed et al., 2019), similar to the Gag-like protein ARC/ARG3.1, which binds RNAs and facilitates intercellular RNA transport among neurons (Ashley et al., 2018; Pastuzyn et al., 2018). Interestingly, the abundance of PEG10-RF1, but not RF1/2, was found to be regulated by the de-ubiquitinase USP9X, which specifically protects the short form against proteasomal degradation (Abed et al., 2019).

CXX1B is expressed from members of the same Ty3/Gypsy retroelement family as PEG10 and is similar to PEG10-RF1 in being composed almost entirely of a Gag-like domain (Supplemental Figure 4C). CXX1B has significant sequence homology to the Gag-like domain of PEG10, as well as the ancestral Gag region of the Sushi-ichi retroelement, though it has little homology to the ancestral Gag region of retroviruses and is lacking the CCHC domain of PEG10 and other Gag-like regions (Supplemental Figure 4D).

Expression of TRIM32, PEG10, and CXX1B is enriched in murine CNS tissues compared to many other tissue types. At the mRNA level, both *Ubqln2* and *Trim32* show high levels of expression in a variety of neuronal subtypes (Supplemental Figure 5A-B). *Peg10* mRNA is most highly enriched in transitional tissues present during development, followed by oligodendrocytes and placental cell types (Supplemental Figure 5C). In contrast to the related *Peg10*, however, *Cxx1b* mRNA is highly expressed in a variety of neuronal subtypes and oligodendrocytes (Supplemental Figure 5D). TMT analysis of multiple murine tissues found enrichment of both UBQLN2 and TRIM32 protein in brains (Supplemental Figure 6A, top), and a similar label-free analysis found PEG10 enrichment in testis and CXX1B enrichment in brain and lung (Supplemental Figure 6A, bottom). Additional studies have observed high levels of PEG10 protein in CNS tissue (Abed et al., 2019; Clark et al., 2007). CXX1B, TRIM32, and UBQLN2 protein were all enriched as least two-fold in brain tissue as compared to liver (Supplemental Figure 6B).

### Ubiquilin-dependence of PEG10, TRIM32, and CXX1B abundance is conserved between mouse and humans

To investigate the mechanism of protein accumulation in *Ubqln*-deficient mouse tissue, and its relevance to humans, a cell culture system was used. WT or *UBQLN1*, *2*, and *4* triple knockout (‘TKO’) HEK 293 cells (Itakura et al., 2016), which are of human origin, were examined for expression of top hits from Figure 2. Cells lacking Ubqlns demonstrated an accumulation of PEG10-RF1/2 but not of PEG10-RF1, as well as accumulation of TRIM32 (Figure 3A). TKO cells also lacked appreciable expression of RTL8, the human ortholog of CXX1B (Figure 3A). In humans, the ortholog of CXX1B is found as three nearly identical proteins which are indistinguishable by Western blot: RTL8A (also known as FAM127B), RTL8B (also known as FAM127C), and RTL8C (known as FAM127A), though murine CXX1B has the highest sequence homology to RTL8C. mRNA levels of each protein were compared between WT and TKO cells. *TRIM32* was unchanged by Ubqln loss, whereas *PEG10* showed modest changes in expression that cannot account for the observed changes at the protein level (Supplemental Figure 7A). In addition, *RTL8* gene expression was upregulated upon Ubqln loss despite a near-total loss of the protein (Supplemental Figure 7A). Importantly, we observed similar mRNA levels of *Peg10*, *Trim32*, and *Cxx1b* in WT versus *Ubqln2^-/-^* murine neuronal tissues (Supplemental Figure 7B-D), indicating that the changes in the level of these proteins were not due to mRNA expression level changes.

**Figure 3:**
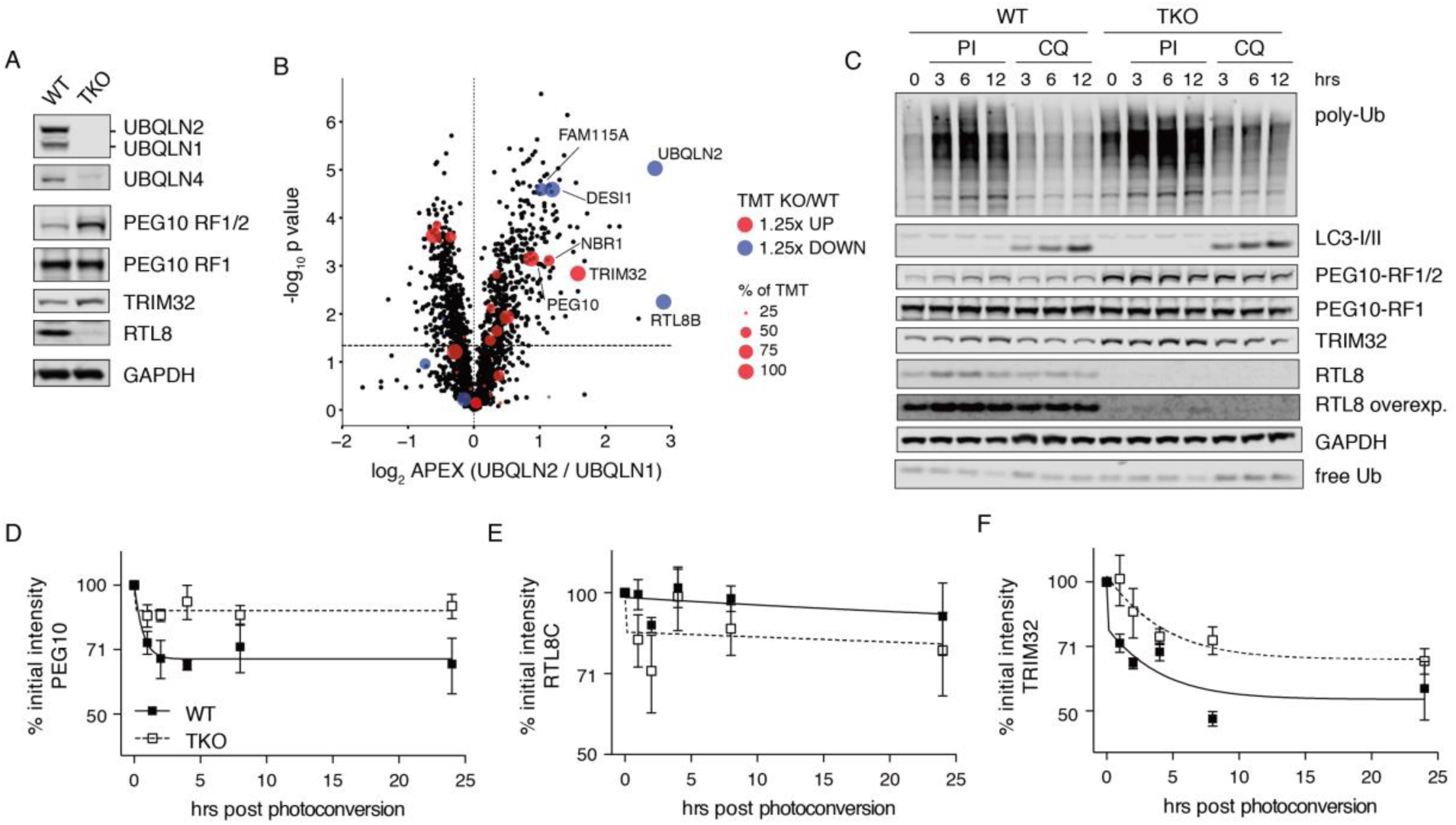
Validation and proteasome dependence of top hits in HEK 293 cells. (A) Western blot demonstrating similar changes to TRIM32, PEG10, and RTL8 protein levels in HEK 293 cells lacking *UBQLN1*, *2*, and *4* (labelled TKO). (B) APEX proximity labeling of TKO HEK 293 cells transfected with plasmids expressing either APEX2-UBQLN1 or APEX2-UBQLN2. Results are depicted as comparison of proteins pulled down more efficiently between conditions. Color of the dots represents the proteins quantified in the APEX study that were altered at least 25% up (red) or 25% down (blue) in *Ubqln2-/-* tissues (KO), with a p value < 0.05, compared to WT littermates (Figure 2). Size of colored dots indicates the number of TMT where each protein was altered at least 25% from all the TMTs where it was quantified (represented as percentages). (C) WT or TKO HEK 293 cells were treated with bortezomib plus epoxomicin (50 nM each), or chloroquine (50 µM), for 0, 3, 6, or 12 hr to inhibit proteasomal or lysosomal degradation respectively. Inhibition of degradation was confirmed with either poly-ubiquitin (Ub) accumulation and free ubiquitin (Ub) depletion, or LC3-I/II conversion respectively. Western blots were probed for top UBQLN2 hits. RTL8 overexp. = overexposure of RTL8 blot. (D-F) WT or TKO HEK 293 cells were transfected with constructs bearing C-terminal fusions of (D) PEG10-RF1/2, (E) RTL8C, and (F) TRIM32 with Dendra2 fluorophore with an IRES-CFP transfection control. After 48 hr, cells were exposed to 60 sec of intense blue light to induce photoconversion of Dendra2 and samples were taken at the indicated timepoints for quantitation of RFP/CFP ratio via flow cytometry. Lines of best fit using a two-phase decay were plotted in Prism for each time course. N=4 independent experiments for each construct.

In addition to examining HEK 293 cells, we generated human ES cells lacking *Ubqln2* and infected with lentivirus encoding doxycycline-inducible Ngn2 expression according to (Zhang et al., 2013). Induced neurons (iNeurons) were tested for protein expression of TRIM32, PEG10, and RTL8. TRIM32 levels were unaltered by *Ubqln2* loss in iNeurons, unlike in mouse tissues (Figure 2) and human HEK cells (Figure 3). PEG10-RF1/2 protein was upregulated between 2- and 4-fold in *Ubqln2^-/-^* cells, whereas PEG10-RF1 was unaffected by *Ubqln2* loss (Supplemental Figure 8A-B). RTL8 protein as measured by Western blot was often completely undetectable in *Ubqln2^-/-^* iNeurons (Supplemental Figure 8A-B), confirming results from mouse and HEK 293 cells. QPCR was performed on mRNA from iNeurons to determine whether these proteins were also regulated at the transcriptional level. mRNA levels were not significantly changed in *Ubqln2^-/-^* iNeurons compared to WT controls (Supplemental Figure 8C). In conclusion, the most promising hits from *Ubqln2^-/-^* proteomics and TKO HEK 293 cells are also dysregulated in human iNeurons lacking functional UBQLN2.

To determine whether TRIM32, PEG10, and CXX1B might bind to UBQLN2, we performed a peroxide-catalyzed proximity biotin-labeling experiment. UBQLN1 and UBQLN2 constructs were fused to a modified plant peroxidase (APEX2) (Lam et al., 2015) at the N-terminus (Supplemental Figure 9A), and the resulting chimeric proteins were transiently transfected into TKO HEK 293 cells (Supplemental Figure 9B). After preincubation in the presence of biotin-phenol, proximity labeling was initiated by the addition of hydrogen peroxide for 1 min, after which the cells were lysed and biotin-labelled proteins pulled down with streptavidin beads. A TMT-10plex analysis was performed on isolated biotinylated proteins in biological triplicate. A direct comparison of the proteins labelled by APEX2-UBQLN2 in comparison to APEX2-UBQLN1 allowed us to confirm RTL8, TRIM32 and PEG10 as some of the most UBQLN2-specific binding partners compared to UBQLN1 (Figure 3B). In addition to previously identified proteins, we also found a number of new proteins which specifically interact with UBQLN2, including FAM115A/TCAF1, which is an interactor of PEG10 (Supplemental Figure 9C and (Huttlin et al., 2017)), NBR1, and DESI1 (Figure 3B). CXX1B, also known as RTL8A-C in humans, also interacts with PEG10 (Supplemental Figure 9C and (Huttlin et al., 2017; Schweppe et al., 2018)), though it is unclear whether this interaction changes the functional state of either protein.

### Proteasome dependence of *Ubqln2* effect

The simplest hypothesis for the accumulation of PEG10 and TRIM32 in *Ubqln2^-/-^* tissues and cells is an impairment of their proteasomal degradation. In support of this hypothesis, ubiquitin-conjugated forms of TRIM32, PEG10, and CXX1B/RTL8 have all previously been observed to accumulate in HTC116 cells treated with proteasome inhibitors (Supplemental Figure 10A) (Rose et al., 2016). To examine their turnover in the context of *Ubqln* deficiency, WT and TKO HEK 293 cells were treated with the proteasome inhibitors or autophagy inhibitors. Inhibition of proteasomal degradation was confirmed by observing elevated levels of ubiquitin-conjugates (Figure 3C); after three hours of treatment, WT cells had accumulated substantial amounts of PEG10-RF1/2 and TRIM32 (Figure 3C). TKO cells did not accumulate appreciable amounts of either protein upon proteasome inhibition, indicating that PEG10-RF1/2 and TRIM32 are indeed degraded through a Ubqln-dependent pathway. Chloroquine treatment had a negligible effect on the accumulation of these proteins despite inhibition of LC3I/II conversion (Figure 3C).

Despite the accumulation of ubiquitin-conjugated CXX1B/RTL8 upon proteasome inhibition of wild-type HTC116 cells (Rose et al. 2016), its levels decreased in *Ubqln2*-deficient cells (Figure 3A), in accord with the mouse data (Figure 2). Ubqlns have occasionally been reported to protect some clients from protein degradation, though evidence for this has largely been limited to cell culture systems involving Ubqln overexpression (Massey et al., 2004). Inhibition of proteasomal degradation resulted in only a minor accumulation of RTL8 protein in TKO cells (Figure 3C and Supplemental Figure 10B), so WT and TKO HEK 293 cells were incubated with high-dose proteasome inhibitors as in Figure 3B (Supplemental Figure 10B). High-dose proteasome inhibition partially rescued TKO RTL8 levels, indicating that its abundance is at least in part regulated by proteasomal degradation.

Since proteasome inhibition did not entirely rescue RTL8 protein levels, we considered whether its synthesis was altered in the absence of *Ubqlns*. To examine the RTL8 synthesis, WT or TKO HEK 293 cells were starved of methionine and incubated with the methionine analog azido-homo-alanine (AHA) to label nascent protein (Supplemental Figure 11A). Following incubation, RTL8 protein was immunoprecipitated. While the total RTL8 protein content in TKO cells is dramatically lower than in WT cells (Supplemental Figure 11A-B), the amount of nascent RTL8 was indistinguishable at 30 min of labeling between the two genotypes (Supplemental Figure 11B), indicating that synthesis of RTL8 is not altered by the loss of Ubqlns.

To confirm the UBQLN-dependence of the degradation of TRIM32, PEG10, and RTL8, we designed a set of fusion proteins with the photo-convertible fluorophore Dendra2 (Gurskaya et al., 2006) at the C-terminus with an IRES-CFP cassette as a control for transfection efficiency (Supplemental Figure 12A). These constructs were expressed in either WT or TKO HEK 293 cells. After 48 hr of expression, the cellular pool of Dendra2-fusion proteins was photoconverted from green (GFP) to red (RFP) by exposure to 488 nm light (Supplemental Figure 12B). Flow cytometric analysis allowed the quantitation of CFP, GFP and photoconverted GFP levels of cells at multiple timepoints following the laser pulse.

PEG10 protein degradation was dramatically delayed in the absence of Ubqln expression (Figure 3D), with very little PEG10 protein degraded in TKO cells during the 24-hr testing period. Results confirming PEG10’s status as a direct client of UBQLN proteins were obtained by WB through cycloheximide chase assays performed on iNeurons expressing endogenous PEG10 protein (Supplemental Figure 12C). Very little RTL8C protein was degraded during the time course in transfected cells, although TKO cells had a modest acceleration in early RTL8C protein loss (Figure 3E), consistent with the hypothesis that UBQLN2 protects CXX1B/RTL8 from degradation. Like PEG10, TRIM32 degradation was also delayed in TKO cells (Figure 3F). Taken together, these results strongly suggest that PEG10 and TRIM32 are bona fide UBQLN2 clients, whereas CXX1B/RTL8C is protected from degradation by UBQLN2.

### Pathway analysis of *Ubqln2^-/-^* tissue reveals *in vivo* neuronal defects in serotonergic signaling

A strength of global proteomic analysis is the ability to identify pathways that link a genetic perturbation to the development of overt phenotypes *in vivo*. To identify families of proteins that were either upregulated or downregulated in *Ubqln2^-/-^* animals, GO-term enrichment was performed on all proteins significantly altered by *Ubqln2* loss. Many pathways were implicated in this proteomic analysis (Supplemental Table 4), the most notable of which are highlighted in Figure 4A. Upregulation of proteins involved in ER stress was evident in aged *Ubqln2^-/-^* brains (Figure 4A) but was not evident in any young tissues (Supplemental Table 4). One cluster of closely related pathways involving neurotransmitter proteins and their transporters was downregulated in both aged and young *Ubqln2^-/-^* brains (Figure 4A). The loss of proteins involved in neurotransmitter synthesis and release was examined more closely: proteins assigned to the GO-term neurotransmitter transporter activity were quantified and compared in aged brains of WT and *Ubqln2^-/-^* littermates for their relative expression (Figure 4B). Among this class of proteins, the serotonin transporter SLC6A4 (also known as SERT or 5-HTT) was one of the most affected (Figure 4B). To confirm alteration of this pathway, spinal cords and brains of WT and *Ubqln2^-/-^* aged animals were immunostained for serotonin, or 5-HT (Figure 4C,E), which was found at significantly lower levels in *Ubqln2^-/-^* tissues as compared to WT counterparts (Figure 4D,F).

**Figure 4:**
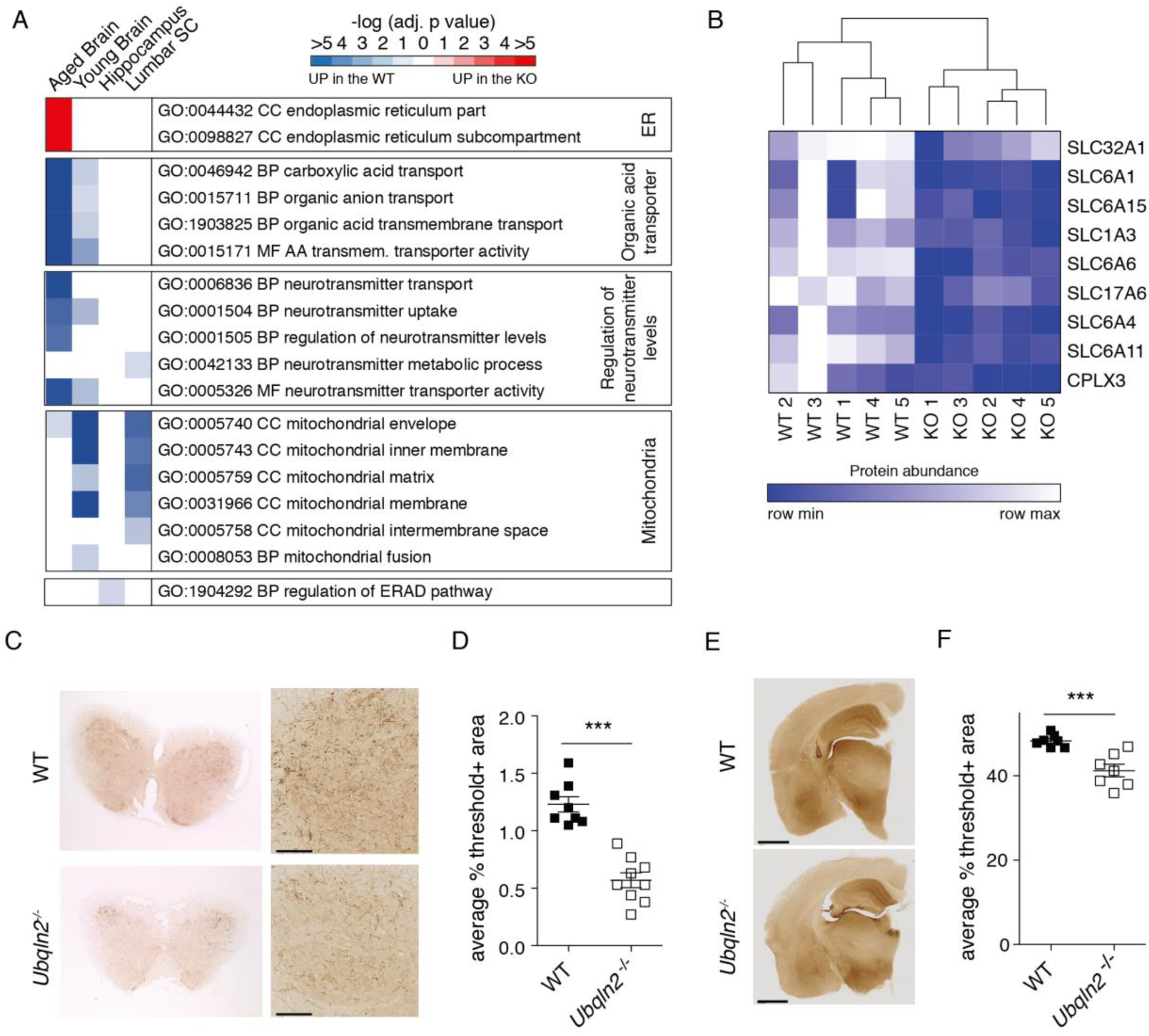
Serotonin staining is decreased in aged *Ubqln2-/-* tissues. (A) GO-term enrichment analysis was performed on brains, hippocampus, and lumbar spinal cord from aged and young mice (from Figure 2) to identify pathways which were altered upon *Ubqln2* loss. Color corresponds to the significance of pathway enrichment as measured by the −log10(adjusted p value) and is either red, for pathways enriched upon *Ubqln2* loss (Up in KO), or blue for pathways downregulated upon *Ubqln2* loss (Up in WT). Notable significantly enriched pathways were selected for visual summarization from Supplemental Table 4. (B) GO-term enrichment of brain hemispheres from aged WT and *Ubqln2-/-* animals (shown as ‘KO’) reveals a decrease in expression of proteins involved in neurotransmitter signaling. Nine significantly altered proteins from the MF GO-term:0005326 neurotransmitter transporter activity were highlighted to demonstrate differences between WT and *Ubqln2-/-* counterparts. Color represents the protein abundance as measured by mass spectrometry, scaled independently for each protein. (C) Lumbar spinal cords from approximately one-year old animals were prepared for histological quantitation of serotonin-positive neuronal projections via 5-HT staining. Shown are representative images of spinal cord cross-section (left) and zoomed in area highlighting neuronal projections. Right scale bar = 100 μm. (D) Quantitation of serotonin positivity as a proportion of area from (C) using thresholding. N=8-9 animals. (E) Coronal sections of brains from young animals were prepared for histological quantitation of serotonin-positive neuronal projections. Scale bar = 1 mm. (F) Quantification of serotonin positivity as a proportion of area using thresholding to determine 5-HT positivity. N=7 animals. Statistical significance for (D,F) was determined using an unpaired, two-tailed Student’s T test.

### Proteomics of ALS disease models with *Ubqln2* mutation

ALS-linked point mutations of *UBQLN2* could in principle be nullimorphic, hypomorphic, hypermorphic, or neomorphic. To assess this at the whole-proteome level, we analyzed isolated hippocampus and spinal cord from a transgenic Ubqln2 animal that overexpresses the P497S allele of human *UBQLN2* (Le et al., 2016). Tissues were taken at a young age (Supplemental Table 2) to minimize downstream effects of late-stage disease, and to facilitate the identification of client proteins before they are obscured by stress responses associated with disease progression. First, we used a proteomic approach to quantify the amount of total UBQLN2 in WT versus transgenic animals: by selecting peptides for examination that are identical between mouse and human UBQLN2, we found that hippocampal tissue of WT Tg mice has approximately twice as much UBQLN2 protein as control animals, and that the degree of overexpression was statistically indistinguishable between animals expressing mutant and WT transgenes (Supplemental Figure 13A).

As a control for effects of overexpression, we initially compared hippocampus and spinal cord proteomes of transgenic mice overexpressing wild-type human UBQLN2 to nontransgenic animals (Suppl. Figure 13B-C). Numerous proteomic hits were found, which is unexpected, given the modest degree of overexpression. Thus, UBQLN2 function appears to be highly sensitive to its levels. The transgene reduced TRIM32 levels in both hippocampal and spinal cord samples, further supporting the view that TRIM32 is a UBQLN2 client. CXX1B levels were elevated upon overexpression of *Ubqln2*, consistent with the hypothesis that UBQLN2 protects CXX1B from degradation, as suggested above. When the nontransgenic mouse, the wild-type transgenic, and the mutant transgenic were then subjected to a pooled TMT analysis, we observed distinct clustering of biological triplicates in both hippocampal and spinal cord samples (Supplemental Figure 13D, E). Results from the transgenic animals (Figure 5A-C) showed an elevation of TRIM32 as a result of the P497S mutation, suggesting that this mutation compromises shuttling of TRIM32 to the proteasome for degradation. Surprisingly this was not the case for PEG10, which was reduced in level by the mutation. Thus, the mutational effects are client protein specific in the context of this transgenic model. Overexpression of the mutant allele of *Ubqln2* also caused enrichment of SQSTM1/P62 in both hippocampus and spinal cord (Figure 5E-F), which along with TRIM32 upregulation in hippocampus was confirmed by Western blot (Supplemental Figure 13F-G).

**Figure 5:**
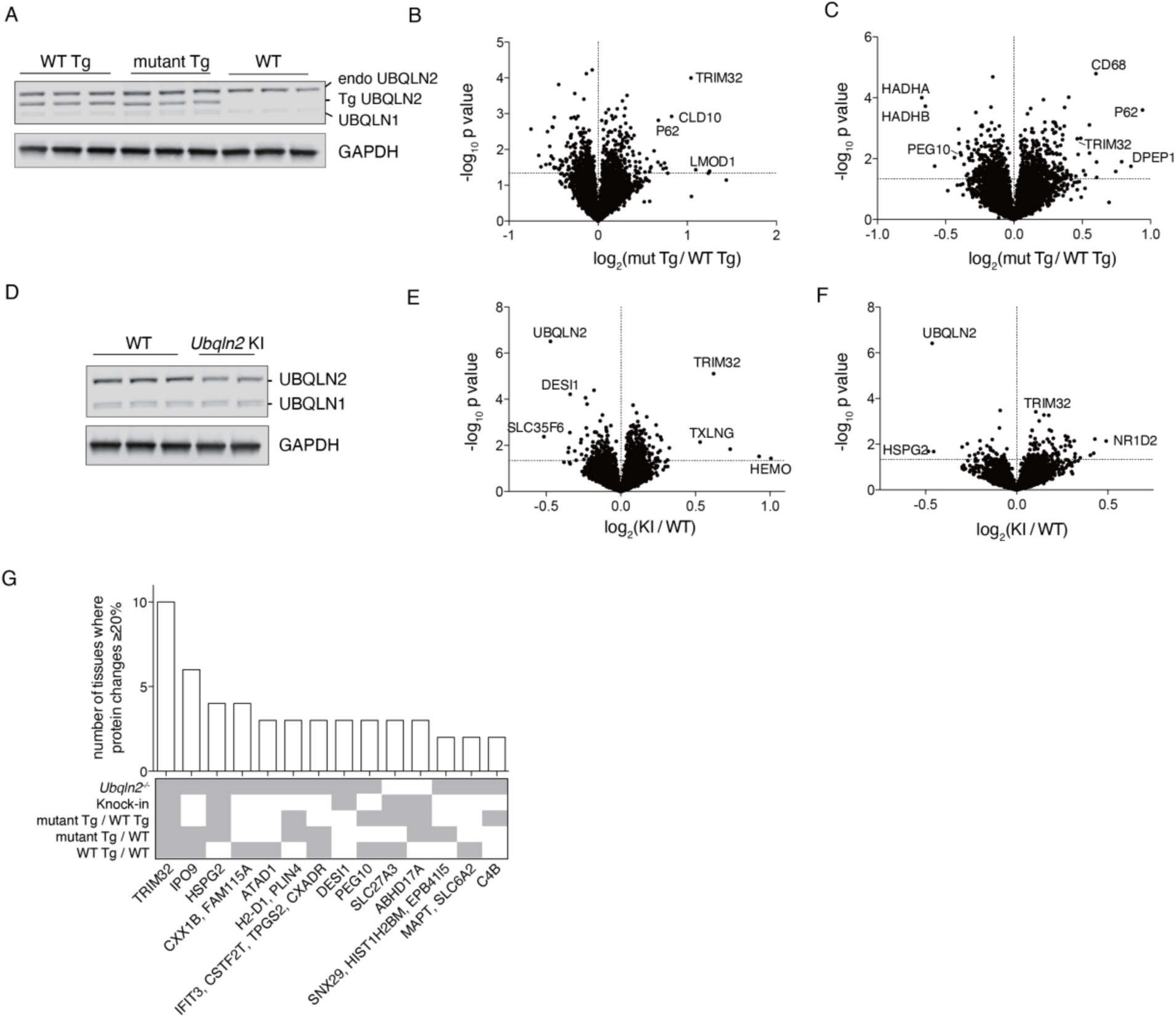
Proteomic meta-analysis of multiple *Ubqln2*-mediated mouse models of ALS. (A) Western blot of endogenous UBQLN2 (‘endo’, top band), human UBQLN2 transgene (‘Tg’, middle band), and UBQLN1 protein (bottom band) in hippocampal tissue of WT transgenic (WT Tg), mutant (P497S) Tg and non-transgenic (WT) mice. GAPDH was used as loading control. (B-C) Hippocampus (B) and lumbar spinal cord (C) from same animals used in (A) were processed for TMT analysis as in Figure 2. N=3 animals per genotype. (D) Western blot of UBQLN2 and UBQLN1 in hippocampus of WT and *Ubqln2* knock-in KI (P506T) mice. Hippocampus (E) and spinal cord (F) of WT or mutant *Ubqln2* KI mice. N= 3 WT animals, 2 KI animals. (G) Individual proteins upregulated or downregulated at least 20%, allowing for standard error, in multiple tissues and/or model systems. The y-axis shows the number of tissues in which the protein is altered, and the x-axis highlights the names of those proteins changed. Below the bar graph is a graphical summary of the genetic backgrounds in which the protein is altered.

We proceeded to examine another characterized mouse model (Hjerpe et al., 2016) which is based on a knock-in (KI) rather than a transgene. The proteomic impact of the P506T knock-in mutant was weaker than that of the transgenic mutant or the null, perhaps due in part to a partial reduction in UBQLN2 level caused by the mutation (Figure 5D-F). However, the TRIM32 effect was still evident in these samples. Other than TRIM32, the proteomic hits were largely distinct when the transgenic and KI models were compared (Figure 5).

### Meta-analysis of proteomic datasets

Individual proteins were assessed for altered levels in all tested *Ubqln2*-mediated models of neurodegenerative disease. Proteins that were altered at least 20% in at least two tissues examined are highlighted in Figure 5G. TRIM32 was the most widely shared protein, with at least 20% alteration in all tissues tested, with representation in each genetic model examined. The nuclear importer IPO9 was shared by 6 tissues among *Ubqln2^-/-^* animals as well as *Ubqln2*-transgenic model systems. The retroelement proteins CXX1B and PEG10 were altered in *Ubqln2* transgenic mouse tissues in addition to *Ubqln2^-/-^* tissue (Figure 5G, Supplemental Figure 13C-D). Thus, interestingly, some features of the null mutant were observed upon overexpression of the WT protein.

In the second form of meta-analysis, a similar GO-term enrichment to that in Figure 4 was performed on hippocampal and spinal cord tissue using proteins significantly altered between WT and mutant conditions (Figure 6A). Metabolic pathways were downregulated in *Ubqln2* mutant conditions, some of which reflected subtle changes to proteins involved in the generation of molecules involved in serotonergic and other neurotransmitter signaling (Supplemental Table 4). However, there was also downregulation of pathways enriched for mitochondrial proteins, especially upon overexpression of mutant UBQLN2 protein (Figure 6A). Mutant UBQLN2 protein expression also caused upregulation of proteasomal degradation pathways, including many proteasome components as well as accessory factors such as SQSTM1/P62 (Supplemental Table 4, Figure 5E-G, Figure 6A). Closer investigation of the cellular compartment GO-term pathway ‘Proteasome complex’ revealed upregulation of a large number of proteasome subunit and accessory proteins in the hippocampus of transgenic mice expressing mutant Ubqln2 (Figure 6B). Western blotting of hippocampal tissue from transgenic mice confirmed the accumulation of ubiquitin conjugates (Figure 6C), consistent with (Le et al., 2016), and suggesting that upregulated proteasome components may be a response to a broad defect in proteasome function as suggested by the accumulation of ubiquitinated proteins upon UBQLN2 dysfunction.

**Figure 6:**
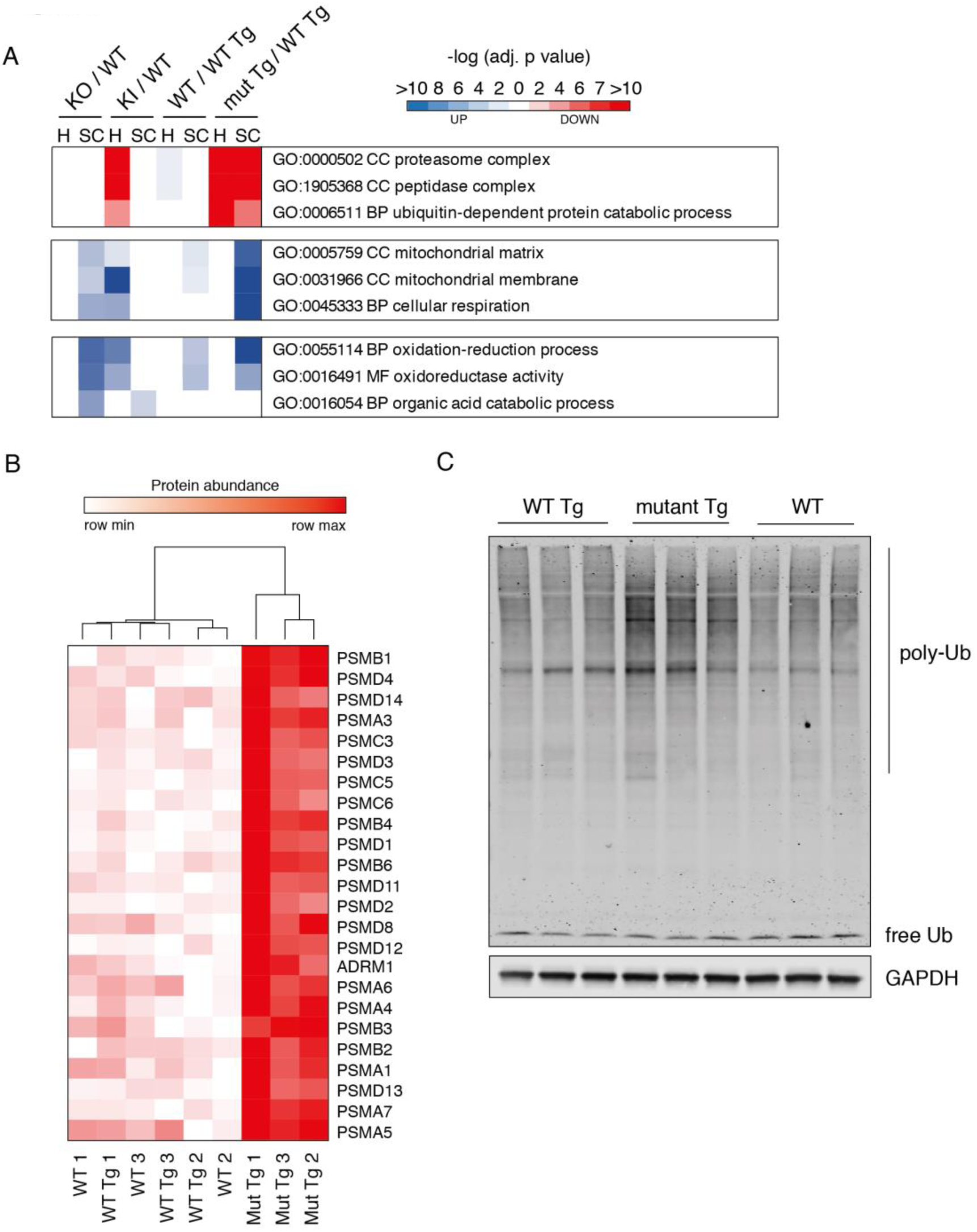
Evidence of alterations in the ubiquitin-proteasome system of *Ubqln2*-Tg animals. (A) GO-term enrichment analysis was performed on hippocampus (H) and lumbar spinal cord (SC) from *Ubqln2-/-*, *Ubqln2* KI (P506T), *Ubqln2* WT transgenic or *Ubqln2* (P497S) mutant transgenic mice compared to their respective controls to identify pathways which were altered upon *Ubqln2* perturbation, as in Figure 4. The most significantly enriched pathways were selected for visual summarization from Supplemental Table 4. (B) GO-term GO:0000502 Cellular Compartment (CC) proteasome complex breakdown of individual proteins in hippocampus from Tg animals. (Similar to Figure 4B). (C) Western blot against ubiquitin (Ub) shows the accumulation of ubiquitin conjugates (poly-Ub) in mutant *Ubqln2*-transgene (mutant Tg) expressing hippocampal tissues of mice. GAPDH was used as loading control.

## Discussion

As Ubqln proteins are substrate delivery factors for the proteasome, the primary working hypothesis for the role of *Ubqln2* in neurodegeneration has been that, upon its perturbation, client protein accumulation leads to the development of disease. However, the physiological clients of UBQLN2 in neuronal cells remained poorly understood and had not been studied in an unbiased, global manner. We performed the first large-scale global proteomic analysis of multiple models of *Ubqln2*-mediated neurodegenerative disease to identify proteins that are under the control of UBQLN2. Quantitative global proteomics offers a view of the mutant or disease phenotype that is exceptional in detail and in scope. Moreover, because we have characterized the *Ubqln2^-/-^* mutant as a baseline, our proteomic data provide strong evidence that the etiological mutations studied here result in both loss-of-function and gain-of-function effects, which interestingly depend on the target protein. For example, since TRIM32 levels were elevated in *Ubqln2* null mutants, reduced in *Ubqln2* overexpressors, and elevated by *Ubqln2* point mutations, stabilization of TRIM32 is not a purely neomorphic phenotype. But while PEG10 is, like TRIM32, elevated in the null, it is, unlike TRIM32, decreased by the P497S mutation. Most likely the P497S mutation attenuates delivery of TRIM32 to the proteasome but does not do so for PEG10.

While previous studies have suggested that Ubqlns can regulate the turnover of a significant fraction of proteasome substrates, the changes to the proteome that we find in disease-relevant tissue *in vivo* are instead focused on a limited number of proteins. Although our proteomics approach is global in scope, it may still miss major clients if for example an explicit stress needs to be imposed to elicit their degradation, or if the mixed cell populations taken from tissues dampen effects that are restricted to specific cell types. Among the strongest hits in our experiments were TRIM32, PEG10, and CXX1B, none of which had previously been identified as UBQLN2 clients. Perturbations of these proteins were found to be mediated by changes in their rates of degradation, at least in part through the proteasome, although this does not exclude the participation of other proteolytic pathways, particularly for CXX1B. Further examination of these hits, via the APEX2 method for example, supported direct interaction between each of these proteins and UBQLN2. To our surprise, however, the strongest hit globally, CXX1B, is actually reduced in level upon loss of UBQLN2 function, indicating that UBQLN2 inhibits rather than promotes CXX1B degradation. The Ubqln effect on CXX1B is very robust as it was observed in murine brain and spinal cord, as well as human HEK 293 cells and induced neurons (known as RTL8 in humans). While the mechanism of CXX1B destabilization remains under investigation, the findings indicate that, depending on the target protein, UBQLN2 is capable of both positive and negative regulation.

A common observation among neurodegenerative conditions is that protein aggregates accumulate at the cellular level well before the onset of overt disease. However, the use of this information for the delay or prevention of disease onset, or even for the development of biomarkers of developing disease, has had limited success. By comparing aged and young animals of the same genetic background, we were able to generate a snapshot of both early and late proteomic changes to relevant disease-related tissues. TRIM32, which was a strong hit in multiple *Ubqln2*-dependent models of disease, was altered even in young brain tissue, as was CXX1B. However, pathway analysis of young brains showed much less involvement of metabolic proteins compared to their aged counterparts. Based on these findings, the increase in TRIM32 and concomitant decrease in CXX1B, both of which were also observed in TKO HEK 293 cells and *Ubqln2^-/-^* iNeurons, may be sensitive early indicators of developing *Ubqln2*-dependent neurodegenerative disease. Further work is underway to see how broadly these changes are found in unrelated cases of sporadic ALS and other neurodegenerative conditions.

In some tissues tested, the most highly upregulated protein upon *Ubqln2* alteration was PEG10; conversely, CXX1B was often the most downregulated protein. Both proteins are members of the Mar retroelement family, which also includes eight other related proteins. Mar retroelements are missing key features of active retrotransposons: they lack necessary elements for transposition in the genome, and its family members are not arranged in tandem; rather, Mar genes are found separated on multiple chromosomes (Brandt et al., 2005). Despite this loss of pre-existing function, multiple members of the Mar family are considered ‘neofunctionalized’. Thus, LDOC1 regulates NF*κ*B activity and sensitizes cells to apoptosis (Inoue et al., 2005; Nagasaki et al., 2003). PEG10 is indispensable for placental development (Abed et al., 2019; Ono et al., 2006) and regulates proliferation (Akamatsu et al., 2015). While the function of CXX1B/RTL8 remains unknown, the multiple genetic duplications that generated RTL8A-C implicate functional diversification (Brandt et al., 2005). Given that UBQLN2 binds to both PEG10 and CXX1B/RTL8, and that PEG10 and CXX1B/RTL8 are similarly binding partners, we speculate that there may be a complex dynamic between the two retroelement proteins that governs their divergent fates upon *Ubqln2* perturbation. Whether the alteration of these two proteins of the Mar retroelement family is involved in the development of *Ubqln2*-dependent neurodegenerative disease is an area of active research.

TRIM32 regulates muscle atrophy through regulated degradation of the proteins dysbindin (Locke et al., 2009) and actin (Cohen et al., 2012). In addition, TRIM32 expression induces neuronal differentiation from precursor cells (Sato et al., 2011). Mutations in the RING domain of TRIM32 cause muscular dystrophy (Kudryashova et al., 2005), but mutations in the BBS domain of the protein cause the developmental disorder Bardet-Biedl Syndrome (Chiang et al., 2006), indicating that the protein has diverse roles in CNS development and function. TRIM32 expression levels at the protein and mRNA levels are elevated upon a variety of CNS insults, including Alzheimer’s disease (Yokota et al., 2006), Duchenne’s muscular dystrophy (Assereto et al., 2016), and acute spinal injury (Liu et al., 2017). In this study, we find a different, post-translational form of TRIM32 regulation mediated by UBQLN2.

By performing pathway analysis on our proteomic results, we were able to identify groups of proteins whose abundance changed upon *Ubqln2* perturbation. One example of this is the downregulation of proteins involved in neurotransmitter signaling evident in both young and aged *Ubqln2^-/-^* brains; none of the proteins in this GO-term were dramatically altered in our proteomic analysis, but a pattern of their alteration emerged upon pathway analysis. Validation of this pathway’s downregulation was performed through histological analysis of brains, which indeed showed decreased 5-HT staining in *Ubqln2^-/-^* tissues. These findings validate the pathway analysis of our proteomics results and provide an intriguing link between protein dysregulation and the emergence of *in vivo* phenotypes. Decreased 5-HT is observed in the plasma, CSF, and spinal cords of ALS patients, and loss of 5-HT neurons leads to spasticity in mouse models of disease, as measured by hind limb clasping (Vermeiren et al., 2018). As such, our proteomic data serve as a potential resource for the discovery of new protein families similarly involved in the ALS disease process.

A second pathway that emerged upon analysis was the upregulation of protein degradation pathways in *Ubqln2*-transgenic animals. The strongest of these hits was P62, which was visually obvious in our volcano plots; alone, this alteration would suggest a highly specific perturbation of selective autophagy. Upon performing pathway analysis, however, there were dozens of proteasome components that were upregulated in transgenic animals, indicating a more generalized dysfunction of protein degradation in these animals. Indeed, these animals have ubiquitin-positive inclusions in neurons of the ventral horn and increased poly-Ubiquitin conjugates in homogenized CNS tissue (Le et al., 2016), which our results confirm.

In conclusion, our study has provided a new understanding of the proteomic landscape of *Ubqln2*-mediated ALS. By leveraging the technology of global proteomics, we have identified proteins in disease-relevant tissues which change upon *Ubqln2* perturbation. Notably, these hits were shared among multiple models of *Ubqln2*-mediated disease, strongly linking them to *Ubqln2* despite fundamental differences in the nature of the genetic perturbation. Furthermore, these hits were also altered in two human models of *Ubqln2* loss: the cell line HEK 293, as well as human ES cells differentiated into neurons. Two of our hits are novel clients of UBQLN2: PEG10 and TRIM32, which accumulate in relevant tissues prior to the development of overt behavioral phenotypes. Future studies will explore the effects of these proteins on neuronal function, and how this relates to RTL8 protein abundance.

## Acknowledgements

The authors thank Dr. Ramanujan Hegde (Medical Research Council, Laboratory for Molecular Biology) for the use of Ubqln TKO HEK cells. The authors thank Dr. Kim Stark for assistance in preparing histological sections of brain and spinal cord for staining and analysis, and Dr. Oded Foreman for pathology support and review. The authors also thank Drs. Jiuchun Zhang and QiaoQiao Wan for assistance with generation of CRISPR hESC cell lines for iNeuron differentiation. This work was supported by a grant to D.F. from the NIH (R01 GM043601) and by a Harvard Brain Science Initiative Seed grant to D.F. A.W. was supported by a grant from the Harvard Medical School Hearst Fund and the Cancer Research Institute Irvington Postdoctoral Fellowship. J.P. was supported by NIH/NIGMS grant R01 GM132129 and S.G. GM97645. T.K. was supported by Motor-Neurone Disease Scotland (MNDS).

## Author Contributions

A.W., M.P., S.dP., A.E., S.G., E.B., and D.F. designed the study. A.W., M.P., S.dP., J.P., M.A., S.D., M.W., H.N., J.S., M.J., T.K., and M.M. conducted experiments. A.W., M.P., S.dP., S.D., M.W., H.N., E.B., and D.F. analyzed the data. A.W., M.P., and D.F. wrote the manuscript.

## Declaration of Interests

S.D., M.W., H.N., A.E., and E.B. are current employees of Genentech, Inc. A.W. is a former employee of Genentech Inc. D.F. is a consultant for Genentech, Inc.

## Methods

### Antibodies and Reagents

Information on antibodies used in the study can be found in Supplemental Table 1.

### Cell lines

HEK 293 cells lacking *UBQLN1*, *2*, and *4* (‘TKO’) were provided by Dr. Ramanujan Hegde (Medical Research Council (MRC) Laboratory of Molecular Biology, Itakura et al., 2016). They were maintained in DMEM (Corning) with penicillin/streptomycin (Gibco), L-glutamine (Gibco), and 10% FBS (Hyclone).

Human ES cells (H9, WiCell Institute) were cultured in fresh E8 medium (G. Chen et al., 2011) on tissue culture plates coated with Matrigel; medium was changed daily and cells were passaged at roughly 50% confluency approximately every three days with 0.5mM EDTA in sterile PBS. SpCas9 protein generated from the expression plasmid pET-NLS-Cas9-6xHis (Addgene #62934) was purified according to (Zuris et al., 2015). sgRNA was generated using the GeneArt Precision gRNA Synthesis Kit (Thermo Fisher Scientific) according to the manufacturer’s instructions and was purified using an RNeasy Mini Kit (Qiagen). To generate *UBQLN2*-deficient ES cells, 0.6 μg of the sgRNA targeting sequence GCCTAAAATCATCAAAGTCA was incubated with 3 μg SpCas9 protein for 10 min at room temperature and electroporated into 2×10^5^ H9 cells. Mutants were identified by Illumina MiSeq and further confirmed by Western blot. Following confirmation of genetic deletion of *UBQLN2*, WT and *UBQLN2^-/-^* ES cells were differentiated into neurons by infecting cells with Ngn2-expressing lentivirus as described (Zhang et al., 2013). After 7 days of differentiation, cells were analyzed for expression of potential UBQLN2 clients.

### Animals

*Ubqln2* knockout animals in the C57BL/6 background were generated by insertion of *loxP* sites flanking the one exon of murine *Ubqln2*. Deletion of *Ubqln2* was induced by transient expression of Cre in ES cells, which were then used to generate live animals. Mice were genotyped using tail clips and the Qiagen DNA extraction kit with FastCycle PCR kit. *Ubqln2^-/-^* mice were housed according to IACUC guidelines and comply with the Institute for Lab Animals’ guidelines for the humane care and use of laboratory animals. All experimental *Ubqln2^-/-^* and WT littermate mice, as well as *Ubqln2* KI and WT littermate mice (Hjerpe et al., 2016), were males to eliminate the confounding variable of gender due to the presence of *Ubqln2* on the X-chromosome (Supplemental Table 2). *Ubqln2* Tg mice (Le et al., 2016) were all heterozygous for transgenic *Ubqln2* allele. For brain and spinal cord tissue collection, mice were anesthetized with 2.5% tribromoethanol (0.5 ml/25 g body weight) and transcardially exsanguinated with phosphate-buffered saline (PBS). Dissected brain and brain hemispheres included intact olfactory lobe, cerebral cortex, and cerebellum. All other euthanasia was performed with CO2 according to IACUC standards, followed by cervical dislocation.

## Behavioral testing

### General

The order of animal testing was randomized by genotype to avoid the potentially confounding effects of recording chambers, arenas, or time of day. Animals were also acclimated to the anteroom of the experimental room (Active avoidance (AA) and fear conditioning (FC) experiments) or the experimental room (all other tests) prior to the beginning of each experiment, and all animals were tested during the day. Between tests, experimental chambers and arenas were cleaned using water or ethanol-containing wipes. To avoid any training sequence effects between AA and FC experiments, the order of AA and FC experiments was also counterbalanced: while half of the mice first underwent FC testing followed by AA testing, the remaining half first underwent AA testing instead. Males used for behavioral testing were housed individually for the duration of all tests and experimenters were blinded to animal genotype for the duration of testing.

### Open Field

An automated PAS Open Field recording system (San Diego Instruments) was used to record spontaneous locomotor activity. Recordings were conducted in a room with bright lights. Mice were placed individually in transparent Plexiglas chambers (40.5 × 40.5 × 38 cm) and horizontal and vertical movements were recorded by two frames in the chamber that were fitted with infrared beams. Activity was measured over 60 min by calculating the total number of beam breaks and rearings per session beginning with the placement of mice into the Plexiglas chambers. Consecutive open field sessions were separated by three days.

### Hind limb clasping

Hind limb clasping was performed by grasping the mouse’s tail near its base and suspending the mouse into the air, clear of all surrounding objects for 20 sec. Scoring was conducted according to (Guyenet et al., 2010).

### Wire hang

Mice were placed on a wire grate and allowed to grasp the wires, after which the grate was gently inverted. The time the mouse was able to hang suspended was recorded, with a maximum latency of 60 sec. Mice unable to reach the full 60 sec could repeat up to 3 times, after which their maximal score was recorded.

### Balance Beam

104 cm-long wooden dowels (of large, medium, and small diameter) were elevated 50 cm above bench surface and angled upwards towards an enclosed escape box. To begin training on balance beam, each mouse was first placed in the escape box for 30 sec, then placed at the start end of the largest balance beam of 24 mm diameter. Mice had 4 consecutive training trials with a maximum acceptable time limit of 1 min to cross. Testing was conducted two days after training and consisted of 9 trials; 3 on the large beam, 3 on the medium (19 mm diameter) and 3 on the small (11 mm diameter). The time required to cross an 80 cm long distance to the escape box on the beam, as well as hind paw foot slips were recorded for each trial.

### Fear conditioning (FC)

FC was conducted according to (Weber et al., 2015). FC chambers for mice (30 × 24 × 24 cm) inside of sound-attenuating cubicles (Med Associates) were used. Each chamber was equipped with a house light, an IR light, a white noise speaker, and a near-IR camera to record movement. The floors of the compartments were equipped with stainless steel metal grids connected to a shock generator. The experiment took two consecutive days: training on day one and a contextual test on the morning of day two followed by a cued test in the afternoon. Furthermore, recording settings of hardware and data acquisition software (Video-Freeze^®^ software, Med Associates) were re-adjusted to account for the different lighting and background conditions for the camera during the cue test. Video data were acquired at a rate of 30 frames per sec; the observation interval was 15 frames or 0.5 sec; and the observation duration was 3 frames or 0.1 sec. The motion threshold was set to 20 (artificial units, AU). Data were acquired with the discrete method: if the motion index was below 20 AU throughout a recording period, freezing was registered. Percent freezing was defined as the percentage of recording periods in which freezing occurred relative to the total number of observation periods. The system was calibrated using camera recordings from empty chambers before the mice were placed inside. FC training began with 3 min of baseline recording, followed by two trials of a 2 sec shock (0.7mA) which served as unconditioned stimulus (UCS) and 30 sec of white noise (∼90 dB; background noise: ∼65 dB) which initially served as the neutral stimulus (NS) or conditioned stimulus (CS). NS or CS co-terminated with the UCS. The two trials were separated by 90s (offset to onset) and the second trial was followed by another 90 sec of recording to measure freezing at the end of the training period. During contextual FC, each animal was re-introduced into the same unaltered chamber where training occurred for a 5 min recording period during which neither white noise nor shock was given. Cued FC testing was conducted in an altered context: for the cue, the house lights were off, the gridded floor was covered with plastic, an insert was placed to alter chamber walls, and the scent of the chamber was changed with a small amount of 1% acetic acid (Mallinckrodt Chemicals). Testing consisted of 3 min of baseline recording, followed by 3 min of white noise without shock. The percent time spent freezing was calculated for each of these experimental phases.

For data analysis, the average motion index across the five 2 sec bins prior to the shocks served as (pre-UCS) baseline relative to the response during the actual 2 sec UCS (during-UCS). The number of fecal boli left in the chamber after the cued-fear condition test period was used as a proxy measure of anxiety and emotionality.

### Active avoidance (AA)

AA experiments were performed according to (Weber et al., 2015) using two-way avoidance chambers inside of sound-attenuating cubicles (Med Associates). Each chamber had two compartments (21 × 16 × 25 cm) that were connected by an auto-guillotine door, and each compartment was equipped with stimulus light and tone generator, as well as four parallel IR light beams located 2.5 cm above the grid floor. Like chambers for FC experiments, the floors of the compartments were connected to a shock generator via stainless steel metal grids. The experimental task was performed according to (Foldi et al., 2011), where each animal received two trainings, consisting of 100 trials each, on two consecutive days.

Sessions began by placing a mouse into one of the two compartments, (selection of which was randomized across the cohort of animals) and letting the mouse acclimatize for 5 min with the door between the compartments closed and all lights off inside the enclosure. Afterwards, each trial began by opening the guillotine door, turning on the stimulus light, and activating the tone (approximately 73 dB). Background noise was approximately 68 dB. If the animal crossed into the second compartment within 5 sec of stimulus: the door was closed, the light and the tone were switched off, an avoidance response was recorded, and the ITI began. If, however, the mouse did not cross within 5 sec, a shock (0.3 mA, 2 sec) was delivered. If the mouse crossed within 2 sec, the shock was stopped, the door was closed, the light and the tone were switched off, an escape response was recorded, and the ITI began. If, however, the mouse did not cross within 2 sec, the shock stopped, the door was closed, the light and the tone were switched off, an escape failure was recorded, and the ITI began. The ITIs ranged from 25 to 55 sec and averaged 40 sec. After the ITI was completed, the next trial was started. This trial-ITI sequence was continued until 100 trials were completed. Hardware and data acquisition were controlled by Med-PC software (Med Associates). Percent avoidance, percent escapes and percent escape failures were calculated using average values obtained from only 5 blocks of 20 consecutive trials, thereby enabling ANOVA at a limited number of factor levels for 100 individual trials.

### Sample preparation for Mass Spectrometry analysis

Tissues were lysed and homogenized in 8 M urea buffer (8 M urea, 150 mM NaCl, 50 mM HEPES (pH 7.5), 75 mM NaCl, 1x EDTA-free protease inhibitor cocktail [Roche], 1x PhosSTOP phosphatase inhibitor cocktail [Roche]). Lysates were centrifuged at 13,000 rpm for 15 min at 4°C and protein concentration quantified by BCA (Pierce). 100 µg of protein of each sample was collected and brought to 1 µg/µl in 8M urea buffer. For reduction and alkylation of cysteines, protein extracts were sequentially incubated with 5 mM TCEP at RT for 30 min, 14 mM iodoacetamide at RT for 30 min in the dark and 10 mM DTT for 15 min at RT. Finally, proteins were methanol-chloroform precipitated and the protein pellet resuspend in 20 mM EPPS (pH 8.5). For digestion, LysC was added at 1:100 (LysC:protein) ratio, and incubated overnight at RT in an orbital shaker at 1,500 rpm. The day after, trypsin was added at 1:75 (trypsin:protein) ratio and incubated for 5 h at 37°C and 1,500 rpm shacking. After digestion, samples were clarified by centrifugation at 13,000 rpm for 10 min and peptide concentration quantified using a quantitative colorimetric peptide assay (Thermo Fisher). For Tandem-Mass-Tag (TMT) labeling we used TMT10-plex kits (Thermo Fisher). In brief, 25 µg of peptides were brought to 1 µg/µl with 200 mM EPPS (pH 8.5), ACN was added to a final concentration of 30% and then 50 µg of each TMT reagent was added (information on all TMT sample labels is included in Supplemental Table 2), followed by incubation of the mixture for 1 hr at RT. Finally, the reaction was stopped by the addition of 0.3% hydroxylamine (Sigma) for 15 min at RT.

To check mixing ratios and labeling efficiency, 1 µg of each sample were polled, desalted, and analyzed by mass spectrometry in a Q-Exactive (Thermo Fisher). Normalization factors calculated from this “label check” (Supplemental Table 5) were used to mix the rest of the samples before desalting with tC18 50 mg SepPak solid-phase extraction cartridges (Waters) and drying using a SpeedVac. Next, dried peptides were resuspended in 10 mM ammonium bicarbonate 5% ACN (pH 8.0) and fractionated with basic-pH reversed-phase high-performance liquid chromatography (using a 3.5μm Zorbax 300 Extended-C18 column [Agilent]). Fractions were collected in a 96-well plate and combined for a total of 24 fractions. 12 of these were desalted following the C18 Stop and Go Extraction Tip (STAGE-Tip) and dried down in the SpeedVac. Finally, peptides were resuspended in 1% formic acid, 3% ACN and analyzed by LC-MS3.

### Liquid chromatography-MS3 (LC-MS3)

All samples were analyzed with an LC-MS3 data collection strategy (McAlister et al., 2014) on Orbitrap Fusion or Orbitrap Fusion Lumos mass spectrometers (Thermo Fisher) equipped with a Proxeon Easy nLC 1000 for online sample handling and peptide separations. Samples resuspended in 1% formic acid, 3% ACN were loaded onto a 100-µm inner diameter fused-silica micro capillary with a needle tip pulled to an internal diameter less than 5 µm. The column was packed in-house to a length of 35 cm with a C18 reverse phase resin (GP118 resin 1.8 μm, 120 Å, Sepax Technologies). Peptides were separated using a 150 min or 135 min linear gradient from 3% to 25% buffer B (100% ACN + 0.125% formic acid) equilibrated with buffer A (3% ACN + 0.125% formic acid) at a flow rate of 400 nL/min across the column. The scan sequence for the mass spectrometer began with an MS1 spectrum, followed with the ‘Top N” (the top 10 precursors) isolation in the quadrupole, CID fragmentation and MS2 detection in the ion trap. Finally, the top ten fragment ion precursors from each MS2 scan were selected for MS3 analysis (synchronous precursor selection, SPS), in which precursors were fragmented by HCD prior to Orbitrap analysis. More information related to the mass spectrometer set up for each TMT are included in Supplemental Table 5.

### TMT-SPS-MS3 data analysis

A suite of in-house software tools was used for .RAW file processing. Searches against a Uniprot mouse or human database (February 2014) were performed using Sequest. Both the forward and reverse sequences, and most common contaminants were included in the database. Database search criteria are as follows: tryptic with two missed cleavages and 2 modifications per peptide, a precursor mass tolerance of 20 ppm, fragment ion mass tolerance of 1.0 Da, static modifications of cysteines (alkylation, 57.02146 Da), and TMT labeling of lysines and N-termini of peptides (+229.162932 Da), and variable oxidation of methionine (15.99491 Da). A controlling peptide and protein level false discovery rates, assembling proteins from peptides, and protein quantification from peptides were applied as previously described (Huttlin et al., 2010; Navarrete-Perea et al., 2018). Each analysis used an SPS-MS3-based TMT method, which has been shown to reduce ion interference compared to MS2 quantification (Paulo et al., 2016). TMT reporter ion intensities were measured using a

0.003 Da window around the theoretical m/z for each reporter ion in the MS3 scan. Peptide spectral matches with poor quality MS3 spectra were excluded from quantitation (<100 summed signal-to-noise across 10 channels and <0.7 precursor isolation specificity). Peptides with a CV < 20 % within each replicate group were used for protein quantification. UBQLN1 and UBQLN2 proteins were quantified using unique peptides. Final protein quantification results were used for pathway analysis. Enrichment of GO-terms (CC, MF, and BP) was performed using the clusterProfiler R package (Yu et al., 2012). For these analyses, all proteins significantly up- or down-regulated between WT and littermate mutant animals were selected. For the GO analysis of each TMT, all proteins quantified in the TMT were used as background.

### Histology and staining

Both the lumbar spinal cord and the left brain hemisphere including intact olfactory lobe, cerebral cortex, and cerebellum, were drop-fixed in 4% PFA for two days at 4°C with agitation and then transferred to PBS for histopathological analyses. Lumbar spinal cord segments and hemi-brains were cryopreserved and were multiply embedded into gelatin matrixes using MultiBrain Technology (NeuroScience Associates, NSA). The lumbar spinal cord block was sectioned coronally at 25 µm and the hemi-brain block was sectioned coronally at 35 µm and sections were sampled according to (Kallop et al., 2014; Le Pichon et al., 2013). Antibody staining was performed at concentrations indicated in Supplemental Table 1. Nissl staining (Fisher Scientific T409-25) was performed according to manufacturer’s recommendations.

### Imaging and quantification of stained sections

Brain and spinal cord tissue samples processed by NSA were imaged on a Leica SCN400 whole slide scanning system (Leica Microsystems) at 200x magnification. Matlab (Mathworks) running on a high performance computing cluster was used for all whole slide image analysis performed in a blinded manner. Motor neuron counts, and quantitation of CD68, GFAP, and Iba1 were performed as previously described (Le Pichon et al., 2013). Analysis of 5-HT staining was performed using color thresholds and morphological operations in Matlab. The percent CD68, GFAP, IBA1 and 5-HT positivity for the entire section was calculated by normalizing the positive pixel area to tissue section area and averaged from eight to twelve sections/animal. All image analysis was performed blind to genotype groups and all images, segmentation overlays, and data were reviewed by a pathologist.

### Metabolic labeling and immunoprecipitation

Confluent cells in 10 cm dishes were starved in methionine free medium (Gibco) for 45 min, then media was replaced with 2 μM azido-homo-alanine (AHA, Life Technologies) in methionine free medium for either 30 min or 4 hr. After incubation, cells were harvested with trypsin, washed, and lysed at a concentration of 2×10^7^/mL of lysis buffer containing 50 mM Tris-HCl (pH 7.4), 150 mM NaCl, and 1% Triton X-100 but no EDTA or *β*-mercaptoethanol. After 30 min on ice, lysate was cleared via centrifugation at 13000 rpm for 10 min at 4°C. 50 μL of lysate was saved for total AHA quantitation and 500 μL of lysate was used for immunoprecipitation of RTL8.

To immunoprecipitate RTL8, 500 μL of lysate plus 500 μL fresh lysis buffer with protease inhibitors was incubated with 7 μg FAM127B antibody (Proteintech) overnight with rocking at 4°C. Following incubation, tubes were spun down and lysate was removed to test for clearance of RTL8. Protein A agarose beads (Invitrogen) were equilibrated with TBS buffer for 2 hr at 4°C with rocking. After one wash, 20 μL of beads were added to each IP tube and rocked for 2 hr at 4°C to bind. After 2 hr, beads were washed 3x with TBS buffer. Then, nascent protein was labelled using standard protocol of the Invitrogen Click-It protein labeling kit with PEG4 biotin-alkyne with 0.5 volumes of buffer. Following labeling but before protein precipitation, tubes were washed 3x with TBS buffer and RTL8 was eluted with 50 μL of 5x Laemmli buffer with BME, then heated for 7 min at 95°C. Total protein from 50 μL of lysate was labelled with PEG4 biotin-alkyne according to manufacturer’s instructions and resuspended in 8M urea buffer.

3 μg of total biotin-labelled protein, from original lysate, as ascertained by BCA (Pierce) of resuspended urea samples, was run on a 4-20% Bis-tris gel with MES buffer (Life Technologies). 30 μL of RTL8 immunoprecipitate was run on a 4-20% Bis-tris gel with MES buffer. Western blots were visualized with Streptavidin IRdye 800 and anti-rabbit IRdye 680 (LICOR).

### Western Blot

For tissue samples and HEK cell lysates, protein was suspended in 8 M urea buffer with 5x Laemmli sample buffer for SDS-PAGE gel. For iNeuron cell samples, protein suspended in Urea buffer was precipitated with methanol/chloroform and resuspended in 2x Laemmli buffer supplemented with *β*-mercaptoethanol and heated for 7 min at 95°C. Samples were run on a 4-20% Tris-glycine gel or 4-12% Nupage Bis-Tris gel (Thermo Fisher) and wet transferred onto PVDF or nitrocellulose paper at 100V for 90 min on ice. Membranes were blocked with Aquablock (East Coast Bio) for at least 20 min and incubated with primary antibody overnight. After three 5-min washes, membranes were incubated for at least 30 min with secondary LICOR antibodies for imaging. After washing again three times for five min each, membranes were visualized and quantitated using LICOR Odyssey imager and software.

### QPCR

For QPCR of brain and spinal cord tissue, frozen samples were homogenized and RNA was extracted using the Qiagen Lipid RNEasy kit with on-column DNA digestion. After RNA extraction, cDNA was generated using the ABI Biosciences One-Step cDNA synthesis kit. QPCR was performed with the following Life Technologies Taqman reagents and Perfecta II mastermix. 10 ng of cDNA was used as input for gene expression assays which were run in triplicate with internal *Gapdh* controls using standard Fast cycling paramaters of an ABI 7500 Fast real-time PCR system. Raw triplicate Ct values were averaged and normalized against *Gapdh*. Taqman primer/probe sets for mouse tissues were as follows: *Gapdh* (VIC): Mm99999915_g1, *Trim32* (MGB): Mm00551733_m1, *Peg10* (MGB): Mm01167724_m1, *Cxx1b* (MGB): Mm03032892_gH.

Primer/probe sets for human iNeurons were as follows: *GAPDH* (VIC):, *TRIM32* (MGB): Hs01045822_m1, *PEG10* (MGB): Hs01122880_m1, *RTL8A* (MGB, also known as FAM127B): Hs00602281_s1, *RTL8B* (MGB, also known as FAM127C): Hs03004887_s1, and *RTL8C* (MGB, also known as FAM127A): Hs05048780_s1.

### Construct design

The human version of the *TRIM32* gene was purchased from Harvard Medical School PlasmID repository. Human full-length *PEG10* encoding both reading frame 1 and reading frame 2, and human *RTL8C*, were cloned from isolated iNeuron cDNA. All three genes were then cloned into LifeAct-Dendra2 from Addgene (plasmid ID: 54694) replacing LifeAct as an N-terminal fusion with Gibson cloning (SGI). Then, an IRES-CFP cassette from pMSCV-IRES-CFP II (pMIC II, Addgene plasmid ID: 52109) was inserted 3’ to the Dendra2 fusion protein, again using Gibson cloning. DH5*α* chemically competent *E. coli* were transformed with expression constructs according to standard protocols and plasmids were purified at 1 ug/μL concentration in sterile, endotoxin-free water using a Zymo midiprep kit. Dendra2 and CFP expression were confirmed by flow cytometry of transfected cells.

The APEX2-*UBQLN1* construct was generated using the *UBQLN1* gene from (Lee et al., 2013). The APEX2-*UBQLN2* construct was generated using cDNA from WT *UBQLN2-*expressing HEK 293 cells (Itakura et al., 2016). Both gene cassettes were inserted into a pCDNA3.1 plasmid containing a V5-APEX2-GS linker cassette using Gibson cloning.

### APEX2 labelling and purification

*UBQLN1*, *2*, and *4* TKO HEK 293 cells were transfected with UBQLN1 or UBQLN2 APEX2 constructs using polyethylenimine (PEI) at a 3:1 PEI:DNA ratio (Polysciences, Inc.). APEX labeling and subsequent streptavidin pull-downs were performed as described in (Paek et al., 2017). Briefly, cells were preincubated in media with 500 µM biotin-phenol (ApexBio) for one hr and then labeled for 1 min by adding hydrogen peroxide to a final concentration of 1 mM. After 1 min, media were aspirated and cells were washed 3x with a quenching buffer containing 5 mM Trolox, 10 mM sodium azide, and 10 mM sodium ascorbate in PBS. Cells were harvested and spun down at 200 *g*, after which the cell pellets were snap-frozen. The cell pellets were lysed in 2 M sodium hydroxide with 7.5% β-mercaptoethanol (Sigma) and proteins were then pelleted after TCA-precipitation. Protein precipitate was reconstituted in 8 M urea buffer containing 100 mM sodium phosphate (pH 8.0), 100 mM ammonium bicarbonate, 1% SDS, and 10 mM TCEP and thoroughly sonicated and vortexed to ensure complete resuspension and TCEP reduction. Samples were spun to remove DNA and supernatant was transferred to new tubes for alkylation with 20 mM iodoacetamide for 25 min. Following alkylation, 50 mM DTT was added to the samples, which were then diluted to 4 M urea and 0.5% SDS. Biotinylated proteins were enriched with streptavidin pulldown using equilibrated streptavidin magnetic beads (Pierce) incubated with protein samples overnight at 4°C. The beads were then washed 3x with wash buffer (4 M urea, 100 mM sodium phosphate [pH 8]) with 0.5% SDS, and 3x with wash buffer without SDS. The beads were subsequently subjected to on-bead digestion with LysC and trypsin. Peptides were then transferred to a new tube, acidified with formic acid (final concentration 3% v/v) and labeled with TMT reagents as previously described. After TMT-labeling, the labelled samples were combined and fractionated using a high pH reversed-phase peptide fractionation kit (Pierce). The resulting 6 fractions and an unfractionated sample were analyzed by LC-MS3 as described above.

### Dendra2 half-life testing

HEK 293 cells were plated to 90% confluency in 12-well plates and transfected with Dendra2 constructs using Lipofectamine 2000 (Life Technologies) and Optimem medium (Gibco). After 24 hr, cells expressing each construct were transferred into 7 wells of flat-bottom 96-well plates for each timepoint of flow cytometry testing. After cells had settled overnight, each well was photoconverted using a fluorescent microscope with 100W halogen bulb. After rapid selection of an optimal field of view, one field of view was photoconverted with continual illumination by blue laser for one min. After one min had elapsed, the next construct was illuminated and photoconverted. Immediately after photoconversion of one timepoint, the samples were rapidly transferred to a round-bottom plate and run on an LSRII flow cytometer (BD Biosciences) with HTS system for data collection. RFP+ cells were gated from CFP+ singlets against background GFP fluorescence, which required compensation. After the RFP+ population was gated, RFP/CFP for the RFP+ population was calculated for each construct at each timepoint and plotted.

### Statistics

For all experiments, statistical significance was measured with either a paired or unpaired Student’s T test (specifics are included in figure legends) with a cutoff of *p<0.05, **p<0.01, ***p<0.001.

## Supplemental Information

### Supplemental Tables

**Supplemental Table 1: Antibodies used in study.**

**Supplemental Table 2: Animals used in study.**

**Supplemental Table 3: Proteomic results from all tissues tested.**

**Supplemental Table 4: Pathway analysis.**

**Supplemental Table 5: TMT label assignment.**

## References Cited

Abed, M., Verschueren, E., Budayeva, H., Liu, P., Kirkpatrick, D.S., Reja, R., Kummerfeld, S.K., Webster, J.D., Gierke, S., Reichelt, M., Anderson, K.R., Newman, R.J., Roose-Girma, M., Modrusan, Z., Pektas, H., Maltepe, E., Newton, K., Dixit, V.M., 2019. The Gag protein PEG10 binds to RNA and regulates trophoblast stem cell lineage specification. PLoS ONE 14, e0214110. doi:10.1371/journal.pone.0214110

Akamatsu, S., Wyatt, A.W., Lin, D., Lysakowski, S., Zhang, F., Kim, S., Tse, C., Wang, K., Mo, F., Haegert, A., Brahmbhatt, S., Bell, R., Adomat, H., Kawai, Y., Xue, H., Dong, X., Fazli, L., Tsai, H., Lotan, T.L., Kossai, M., Mosquera, J.M., Rubin, M.A., Beltran, H., Zoubeidi, A., Wang, Y., Gleave, M.E., Collins, C.C., 2015. The Placental Gene PEG10 Promotes Progression of Neuroendocrine Prostate Cancer. Cell Reports 12, 922–936. doi:10.1016/j.celrep.2015.07.012

Alexander, E.J., Niaki, A.G., Zhang, T., Sarkar, J., Liu, Y., Nirujogi, R.S., Pandey, A., Myong, S., Wang, J., 2018. Ubiquilin 2 modulates ALS/FTD-linked FUS–RNA complex dynamics and stress granule formation. Proceedings of the National Academy of Sciences 115, E11485–E11494. doi:10.1073/pnas.1811997115

Ashley, J., Cordy, B., Lucia, D., Fradkin, L.G., Budnik, V., Thomson, T., 2018. Retrovirus-like Gag Protein Arc1 Binds RNA and Traffics across Synaptic Boutons. Cell 172, 262–274.e11. doi:10.1016/j.cell.2017.12.022

Assereto, S., Piccirillo, R., Baratto, S., Scudieri, P., Fiorillo, C., Massacesi, M., Traverso, M., Galietta, L.J., Bruno, C., Minetti, C., Zara, F., Gazzerro, E., 2016. The ubiquitin ligase tripartite-motif-protein 32 is induced in Duchenne muscular dystrophy. Lab. Invest. 96, 862–871. doi:10.1038/labinvest.2016.63

Ayadi, El, A., Stieren, E.S., Barral, J.M., Boehning, D., 2012. Ubiquilin-1 regulates amyloid precursor protein maturation and degradation by stimulating K63-linked polyubiquitination of lysine 688. Proceedings of the National Academy of Sciences of the United States of America 109, 13416–13421. doi:10.1073/pnas.1206786109

Biggins, S., Ivanovska, I., Rose, M.D., 1996. Yeast ubiquitin-like genes are involved in duplication of the microtubule organizing center. The Journal of Cell Biology 133, 1331–1346. doi:10.1083/jcb.133.6.1331

Blokhuis, A.M., Groen, E.J.N., Koppers, M., van den Berg, L.H., Pasterkamp, R.J., 2013. Protein aggregation in amyotrophic lateral sclerosis. Acta Neuropathologica 125, 777–794. doi:10.1007/s00401-013-1125-6

Brandt, J., Veith, A.M., Volff, J.-N., 2005. A family of neofunctionalized Ty3/gypsy retrotransposon genes in mammalian genomes. Cytogenet. Genome Res. 110, 307– 317. doi:10.1159/000084963

Chang, L., Monteiro, M.J., 2015. Defective Proteasome Delivery of Polyubiquitinated Proteins by Ubiquilin-2 Proteins Containing ALS Mutations. PLoS ONE 10, e0130162. doi:10.1371/journal.pone.0130162

Chen, G., Gulbranson, D.R., Hou, Z., Bolin, J.M., Ruotti, V., Probasco, M.D., Smuga-Otto, K., Howden, S.E., Diol, N.R., Propson, N.E., Wagner, R., Lee, G.O., Antosiewicz-Bourget, J., Teng, J.M.C., Thomson, J.A., 2011. Chemically defined conditions for human iPSC derivation and culture. Nat Meth 8, 424–429. doi:10.1038/nmeth.1593

Chen, X., Ebelle, D.L., Wright, B.J., Sridharan, V., Hooper, E., Walters, K.J., 2019. Structure of hRpn10 Bound to UBQLN2 UBL Illustrates Basis for Complementarity between Shuttle Factors and Substrates at the Proteasome. Journal of Molecular Biology 431, 939–955. doi:10.1016/j.jmb.2019.01.021

Chen, X., Randles, L., Shi, K., Tarasov, S.G., Aihara, H., Walters, K.J., 2016. Structures of Rpn1 T1:Rad23 and hRpn13:hPLIC2 Reveal Distinct Binding Mechanisms between Substrate Receptors and Shuttle Factors of the Proteasome. Structure 24, 1257–1270. doi:10.1016/j.str.2016.05.018

Chen, Y.-C., Umanah, G.K.E., Dephoure, N., Andrabi, S.A., Gygi, S.P., Dawson, T.M., Dawson, V.L., Rutter, J., 2014. Msp1/ATAD1 maintains mitochondrial function by facilitating the degradation of mislocalized tail-anchored proteins. EMBO J 33, 1548– 1564. doi:10.15252/embj.201487943

Chiang, A.P., Beck, J.S., Yen, H.-J., Tayeh, M.K., Scheetz, T.E., Swiderski, R.E., Nishimura, D.Y., Braun, T.A., Kim, K.-Y.A., Huang, J., Elbedour, K., Carmi, R., Slusarski, D.C., Casavant, T.L., Stone, E.M., Sheffield, V.C., 2006. Homozygosity mapping with SNP arrays identifies TRIM32, an E3 ubiquitin ligase, as a Bardet-Biedl syndrome gene (BBS11). Proceedings of the National Academy of Sciences 103, 6287–6292. doi:10.1073/pnas.0600158103

Clark, M.B., Jänicke, M., Gottesbühren, U., Kleffmann, T., Legge, M., Poole, E.S., Tate, W.P., 2007. Mammalian gene PEG10 expresses two reading frames by high efficiency -1 frameshifting in embryonic-associated tissues. Journal of Biological Chemistry 282, 37359–37369. doi:10.1074/jbc.M705676200

Cohen, S., Zhai, B., Gygi, S.P., Goldberg, A.L., 2012. Ubiquitylation by Trim32 causes coupled loss of desmin, Z-bands, and thin filaments in muscle atrophy. The Journal of Cell Biology 198, 575–589. doi:10.1083/jcb.201110067

Dao, T.P., Kolaitis, R.-M., Kim, H.J., O’Donovan, K., Martyniak, B., Colicino, E., Hehnly, H., Taylor, J.P., Castañeda, C.A., 2018. Ubiquitin Modulates Liquid-Liquid Phase Separation of UBQLN2 via Disruption of Multivalent Interactions. Molecular Cell 69, 965–978.e6. doi:10.1016/j.molcel.2018.02.004

Daoud, H., Suhail, H., Szuto, A., Camu, W., Salachas, F., Meininger, V., Bouchard, J.-P., Dupré, N., Dion, P.A., Rouleau, G.A., 2012. UBQLN2 mutations are rare in French and French-Canadian amyotrophic lateral sclerosis. Neurobiol. Aging 33, 2230.e1– 2230.e5. doi:10.1016/j.neurobiolaging.2012.03.015

Deng, H.-X., Chen, W., Hong, S.-T., Boycott, K.M., Gorrie, G.H., Siddique, N., Yang, Y., Fecto, F., Shi, Y., Zhai, H., Jiang, H., Hirano, M., Rampersaud, E., Jansen, G.H., Donkervoort, S., Bigio, E.H., Brooks, B.R., Ajroud, K., Sufit, R.L., Haines, J.L., Mugnaini, E., Pericak-Vance, M.A., Siddique, T., 2011. Mutations in UBQLN2 cause dominant X-linked juvenile and adult-onset ALS and ALS/dementia. Nature 477, 211– 215. doi:10.1038/nature10353

Elsasser, S., Finley, D., 2005. Delivery of ubiquitinated substrates to protein-unfolding machines. Nat. Cell Biol. 7, 742–749. doi:10.1038/ncb0805-742

Elsasser, S., Gali, R.R., Schwickart, M., Larsen, C.N., Leggett, D.S., Müller, B., Feng, M.T., Tübing, F., Dittmar, G.A.G., Finley, D., 2002. Proteasome subunit Rpn1 binds ubiquitin-like protein domains. Nat. Cell Biol. 4, 725–730. doi:10.1038/ncb845

Fecto, F., Yan, J., Vemula, S.P., Liu, E., Yang, Y., Chen, W., Zheng, J.G., Shi, Y., Siddique, N., Arrat, H., Donkervoort, S., Ajroud-Driss, S., Sufit, R.L., Heller, S.L., Deng, H.-X., Siddique, T., 2011. SQSTM1 mutations in familial and sporadic amyotrophic lateral sclerosis. Arch. Neurol. 68, 1440–1446. doi:10.1001/archneurol.2011.250

Finley, D., 2009. Recognition and Processing of Ubiquitin-Protein Conjugates by the Proteasome. Annu. Rev. Biochem. 78, 477–513. doi:10.1146/annurev.biochem.78.081507.101607

Foldi, C.J., Eyles, D.W., McGrath, J.J., Burne, T.H.J., 2011. The effects of breeding protocol in C57BL/6J mice on adult offspring behaviour. PLoS ONE 6, e18152. doi:10.1371/journal.pone.0018152

Ford, D.L., Monteiro, M.J., 2006. Dimerization of ubiquilin is dependent upon the central region of the protein: evidence that the monomer, but not the dimer, is involved in binding presenilins. Biochem. J. 399, 397–404. doi:10.1042/BJ20060441

Funakoshi, M., Sasaki, T., Nishimoto, T., Kobayashi, H., 2002. Budding yeast Dsk2p is a polyubiquitin-binding protein that can interact with the proteasome. Proceedings of the National Academy of Sciences 99, 745–750. doi:10.1073/pnas.012585199

Gilpin, K.M., Chang, L., Monteiro, M.J., 2015. ALS-linked mutations in ubiquilin-2 or hnRNPA1 reduce interaction between ubiquilin-2 and hnRNPA1. Human Molecular Genetics 24, 2565–2577. doi:10.1093/hmg/ddv020

Gurskaya, N.G., Verkhusha, V.V., Shcheglov, A.S., Staroverov, D.B., Chepurnykh, T.V., Fradkov, A.F., Lukyanov, S., Lukyanov, K.A., 2006. Engineering of a monomeric green-to-red photoactivatable fluorescent protein induced by blue light. Nat. Biotechnol. 24, 461–465. doi:10.1038/nbt1191

Guyenet, S.J., Furrer, S.A., Damian, V.M., Baughan, T.D., La Spada, A.R., Garden, G.A., 2010. A simple composite phenotype scoring system for evaluating mouse models of cerebellar ataxia. JoVE e1787. doi:10.3791/1787

Harman, C.A., Monteiro, M.J., 2019. The specificity of ubiquitin binding to ubiquilin-1 is regulated by sequences besides its UBA domain. Biochim Biophys Acta Gen Subj. doi:10.1016/j.bbagen.2019.06.002

Hjerpe, R., Aillet, F., Lopitz-Otsoa, F., Lang, V., England, P., Rodriguez, M.S., 2009. Efficient protection and isolation of ubiquitylated proteins using tandem ubiquitin-binding entities. EMBO Reports 10, 1250–1258. doi:10.1038/embor.2009.192

Hjerpe, R., Bett, J.S., Keuss, M.J., Solovyova, A., McWilliams, T.G., Johnson, C., Sahu, I., Varghese, J., Wood, N., Wightman, M., Osborne, G., Bates, G.P., Glickman, M.H., Trost, M., Knebel, A., Marchesi, F., Kurz, T., 2016. UBQLN2 Mediates Autophagy-Independent Protein Aggregate Clearance by the Proteasome. Cell 166, 1–15. doi:10.1016/j.cell.2016.07.001

Hruz, T., Laule, O., Szabo, G., Wessendorp, F., Bleuler, S., Oertle, L., Widmayer, P., Gruissem, W., Zimmermann, P., 2008. Genevestigator v3: a reference expression database for the meta-analysis of transcriptomes. Adv Bioinformatics 2008, 420747– 5. doi:10.1155/2008/420747

Huttlin, E.L., Bruckner, R.J., Paulo, J.A., Cannon, J.R., Ting, L., Baltier, K., Colby, G., Gebreab, F., Gygi, M.P., Parzen, H., Szpyt, J., Tam, S., Zarraga, G., Pontano-Vaites, L., Swarup, S., White, A.E., Schweppe, D.K., Rad, R., Erickson, B.K., Obar, R.A., Guruharsha, K.G., Li, K., Artavanis-Tsakonas, S., Gygi, S.P., Harper, J.W., 2017. Architecture of the human interactome defines protein communities and disease networks. Nature 545, 505–509. doi:10.1038/nature22366

Huttlin, E.L., Jedrychowski, M.P., Elias, J.E., Goswami, T., Rad, R., Beausoleil, S.A., Villén, J., Haas, W., Sowa, M.E., Gygi, S.P., 2010. A Tissue-Specific Atlas of Mouse Protein Phosphorylation and Expression. Cell 143, 1174–1189. doi:10.1016/j.cell.2010.12.001

Inoue, M., Takahashi, K., Niide, O., Shibata, M., Fukuzawa, M., Ra, C., 2005. LDOC1, a novel MZF-1-interacting protein, induces apoptosis. FEBS Letters 579, 604–608. doi:10.1016/j.febslet.2004.12.030

Itakura, E., Zavodszky, E., Shao, S., Wohlever, M.L., Keenan, R.J., Hegde, R.S., 2016. Ubiquilins Chaperone and Triage Mitochondrial Membrane Proteins for Degradation. Molecular Cell 63, 21–33. doi:10.1016/j.molcel.2016.05.020

Kallop, D.Y., Meilandt, W.J., Gogineni, A., Easley-Neal, C., Wu, T., Jubb, A.M., Yaylaoglu, M., Shamloo, M., Tessier-Lavigne, M., Scearce-Levie, K., Weimer, R.M., 2014. A death receptor 6-amyloid precursor protein pathway regulates synapse density in the mature CNS but does not contribute to Alzheimer’s disease-related pathophysiology in murine models. Journal of Neuroscience 34, 6425–6437. doi:10.1523/JNEUROSCI.4963-13.2014

Kiernan, M.C., Vucic, S., Cheah, B.C., Turner, M.R., Eisen, A., Hardiman, O., Burrell, J.R., Zoing, M.C., 2011. Amyotrophic lateral sclerosis. The Lancet 377, 942–955. doi:10.1016/S0140-6736(10)61156-7

Kleijnen, M.F., Alarcon, R.M., Howley, P.M., 2003. The ubiquitin-associated domain of hPLIC-2 interacts with the proteasome. Molecular Biology of the Cell 14, 3868–3875. doi:10.1091/mbc.e02-11-0766

Kleijnen, M.F., Shih, A.H., Zhou, P., Kumar, S., Soccio, R.E., 2000. The hPLIC proteins may provide a link between the ubiquitination machinery and the proteasome. Molecular Cell 6, 408–419. doi:10.1016/S1097-2765(00)00040-X

Kudryashova, E., Kudryashov, D., Kramerova, I., Spencer, M.J., 2005. Trim32 is a ubiquitin ligase mutated in limb girdle muscular dystrophy type 2H that binds to skeletal muscle myosin and ubiquitinates actin. Journal of Molecular Biology 354, 413–424. doi:10.1016/j.jmb.2005.09.068

Lam, S.S., Martell, J.D., Kamer, K.J., Deerinck, T.J., Ellisman, M.H., Mootha, V.K., Ting, A.Y., 2015. Directed evolution of APEX2 for electron microscopy and proximity labeling. Nat Meth 12, 51–54. doi:10.1038/nmeth.3179

Le Pichon, C.E., Dominguez, S.L., Solanoy, H., Ngu, H., Lewin-Koh, N., Chen, M., Eastham-Anderson, J., Watts, R., Scearce-Levie, K., 2013. EGFR Inhibitor Erlotinib Delays Disease Progression but Does Not Extend Survival in the SOD1 Mouse Model of ALS. PLoS ONE 8, e62342. doi:10.1371/journal.pone.0062342

Le, N.T.T., Chang, L., Kovlyagina, I., Georgiou, P., Safren, N., Braunstein, K.E., Kvarta, M.D., Van Dyke, A.M., LeGates, T.A., Philips, T., Morrison, B.M., Thompson, S.M., Puche, A.C., Gould, T.D., Rothstein, J.D., Wong, P.C., Monteiro, M.J., 2016. Motor neuron disease, TDP-43 pathology, and memory deficits in mice expressing ALS-FTD-linked UBQLN2 mutations. Proceedings of the National Academy of Sciences of the United States of America 113, E7580–E7589. doi:10.1073/pnas.1608432113

Lee, D.Y., Arnott, D., Brown, E.J., 2013. Ubiquilin4 is an adaptor protein that recruits Ubiquilin1 to the autophagy machinery. EMBO Reports 14, 373–381. doi:10.1038/embor.2013.22

Liu, C., van Dyk, D., Li, Y., Andrews, B., Rao, H., 2009. A genome-wide synthetic dosage lethality screen reveals multiple pathways that require the functioning of ubiquitin-binding proteins Rad23 and Dsk2. BMC Biology 7, 75. doi:10.1186/1741-7007-7-75

Liu, Y., Wu, W., Yang, H., Zhou, Z., Zhu, X., Sun, C., Liu, Y., Yu, Z., Chen, Y., Wang, Y., 2017. Upregulated Expression of TRIM32 Is Involved in Schwann Cell Differentiation, Migration and Neurite Outgrowth After Sciatic Nerve Crush. Neurochem. Res. 42, 1084–1095. doi:10.1007/s11064-016-2142-3

Locke, M., Tinsley, C.L., Benson, M.A., Blake, D.J., 2009. TRIM32 is an E3 ubiquitin ligase for dysbindin. Human Molecular Genetics 18, 2344–2358. doi:10.1093/hmg/ddp167

Lomash, R.M., Gu, X., Youle, R.J., Lu, W., Roche, K.W., 2015. Neurolastin, a Dynamin Family GTPase, Regulates Excitatory Synapses and Spine Density. Cell Reports 12, 743–751. doi:10.1016/j.celrep.2015.06.064

Mah, A.L., Perry, G., Smith, M.A., Monteiro, M.J., 2000. Identification of ubiquilin, a novel presenilin interactor that increases presenilin protein accumulation. The Journal of Cell Biology 151, 847–862. doi:10.1083/jcb.151.4.847

Marín, I., 2014. The ubiquilin gene family: evolutionary patterns and functional insights. BMC Evol. Biol. 14, 63. doi:10.1186/1471-2148-14-63

Maruyama, H., Morino, H., Ito, H., Izumi, Y., Kato, H., Watanabe, Y., Kinoshita, Y., Kamada, M., Nodera, H., Suzuki, H., Komure, O., Matsuura, S., Kobatake, K., Morimoto, N., Abe, K., Suzuki, N., Aoki, M., Kawata, A., Hirai, T., Kato, T., Ogasawara, K., Hirano, A., Takumi, T., Kusaka, H., Hagiwara, K., Kaji, R., Kawakami, H., 2010. Mutations of optineurin in amyotrophic lateral sclerosis. Nature 465, 223–226. doi:10.1038/nature08971

Massey, L.K., Mah, A.L., Ford, D.L., Miller, J., Liang, J., Doong, H., Monteiro, M.J., 2004. Overexpression of ubiquilin decreases ubiquitination and degradation of presenilin proteins. J. Alzheimers Dis. 6, 79–92. doi:10.3233/jad-2004-6109

McAlister, G.C., Nusinow, D.P., Jedrychowski, M.P., Wühr, M., Huttlin, E.L., Erickson, B.K., Rad, R., Haas, W., Gygi, S.P., 2014. MultiNotch MS3 Enables Accurate, Sensitive, and Multiplexed Detection of Differential Expression across Cancer Cell Line Proteomes. Anal. Chem. 86, 7150–7158. doi:10.1021/ac502040v

Medicherla, B., Kostova, Z., Schaefer, A., Wolf, D.H., 2004. A genomic screen identifies Dsk2p and Rad23p as essential components of ER-associated degradation. EMBO Reports 5, 692–697. doi:10.1038/sj.embor.7400164

Nagasaki, K., Schem, C., Kaisenberg, von, C., Biallek, M., Rösel, F., Jonat, W., Maass, N., 2003. Leucine-zipper protein, LDOC1, inhibits NF-kappaB activation and sensitizes pancreatic cancer cells to apoptosis. International Journal of Cancer 105, 454–458. doi:10.1002/ijc.11122

Navarrete-Perea, J., Yu, Q., Gygi, S.P., Paulo, J.A., 2018. Streamlined Tandem Mass Tag (SL-TMT) Protocol: An Efficient Strategy for Quantitative (Phospho)proteome Profiling Using Tandem Mass Tag-Synchronous Precursor Selection-MS3. J. Proteome Res. 17, 2226–2236. doi:10.1021/acs.jproteome.8b00217

N’Diaye, E.-N., Debnath, J., Brown, E.J., 2009a. Ubiquilins accelerate autophagosome maturation and promote cell survival during nutrient starvation. autophagy 5, 573– 575.

N’Diaye, E.-N., Kajihara, K.K., Hsieh, I., Morisaki, H., Debnath, J., Brown, E.J., 2009b. PLIC proteins or ubiquilins regulate autophagy-dependent cell survival during nutrient starvation. EMBO Reports 10, 173–179. doi:10.1038/embor.2008.238

Okreglak, V., Walter, P., 2014. The conserved AAA-ATPase Msp1 confers organelle specificity to tail-anchored proteins. Proceedings of the National Academy of Sciences of the United States of America 111, 8019–8024. doi:10.1073/pnas.1405755111

Ono, R., Nakamura, K., Inoue, K., Naruse, M., Usami, T., Wakisaka-Saito, N., Hino, T., Suzuki-Migishima, R., Ogonuki, N., Miki, H., Kohda, T., Ogura, A., Yokoyama, M., Kaneko-Ishino, T., Ishino, F., 2006. Deletion of Peg10, an imprinted gene acquired from a retrotransposon, causes early embryonic lethality. Nat Genet 38, 101–106. doi:10.1038/ng1699

Paek, J., Kalocsay, M., Staus, D.P., Wingler, L., Pascolutti, R., Paulo, J.A., Gygi, S.P., Kruse, A.C., 2017. Multidimensional Tracking of GPCR Signaling via Peroxidase-Catalyzed Proximity Labeling. Cell 169, 338–349.e11. doi:10.1016/j.cell.2017.03.028

Pastuzyn, E.D., Day, C.E., Kearns, R.B., Kyrke-Smith, M., Taibi, A.V., McCormick, J., Yoder, N., Belnap, D.M., Erlendsson, S., Morado, D.R., Briggs, J.A.G., Feschotte, C., Shepherd, J.D., 2018. The Neuronal Gene Arc Encodes a Repurposed Retrotransposon Gag Protein that Mediates Intercellular RNA Transfer. Cell 172, 275–288.e18. doi:10.1016/j.cell.2017.12.024

Paulo, J.A., O’Connell, J.D., Gygi, S.P., 2016. A Triple Knockout (TKO) Proteomics Standard for Diagnosing Ion Interference in Isobaric Labeling Experiments. J. Am. Soc. Mass Spectrom. 27, 1620–1625. doi:10.1007/s13361-016-1434-9

Rao, H., Sastry, A., 2002. Recognition of specific ubiquitin conjugates is important for the proteolytic functions of the ubiquitin-associated domain proteins Dsk2 and Rad23. Journal of Biological Chemistry 277, 11691–11695. doi:10.1074/jbc.M200245200

Rose, C.M., Isasa, M., Ordureau, A., Prado, M.A., Beausoleil, S.A., Jedrychowski, M.P., Finley, D.J., Harper, J.W., Gygi, S.P., 2016. Highly Multiplexed Quantitative Mass Spectrometry Analysis of Ubiquitylomes. Cell Syst 3, 395–403.e4. doi:10.1016/j.cels.2016.08.009

Ross, C.A., Poirier, M.A., 2004. Protein aggregation and neurodegenerative disease. Nat Med 10, S10–S17. doi:10.1038/nm1066

Rothenberg, C., Srinivasan, D., Mah, L., Kaushik, S., Peterhoff, C.M., Ugolino, J., Fang, S., Cuervo, A.M., Nixon, R.A., Monteiro, M.J., 2010. Ubiquilin functions in autophagy and is degraded by chaperone-mediated autophagy. Human Molecular Genetics 19, 3219–3232. doi:10.1093/hmg/ddq231

Rutherford, N.J., Lewis, J., Clippinger, A.K., Thomas, M.A., Adamson, J., Cruz, P.E., Cannon, A., Xu, G., Golde, T.E., Shaw, G., Borchelt, D.R., Giasson, B.I., 2013. Unbiased Screen Reveals Ubiquilin-1 and -2 Highly Associated with Huntingtin Inclusions. Brain Research 1524, 62–73. doi:10.1016/j.brainres.2013.06.006

Saeki, Y., Saitoh, A., Toh-e, A., Yokosawa, H., 2002. Ubiquitin-like proteins and Rpn10 play cooperative roles in ubiquitin-dependent proteolysis. Biochemical and Biophysical Research Communications 293, 986–992. doi:10.1016/S0006-291X(02)00340-6

Sato, T., Okumura, F., Kano, S., Kondo, T., Ariga, T., Hatakeyama, S., 2011. TRIM32 promotes neural differentiation through retinoic acid receptor-mediated transcription. Journal of Cell Science 124, 3492–3502. doi:10.1242/jcs.088799

Schweppe, D.K., Huttlin, E.L., Harper, J.W., Gygi, S.P., 2018. BioPlex Display: An Interactive Suite for Large-Scale AP-MS Protein-Protein Interaction Data. J. Proteome Res. 17, 722–726. doi:10.1021/acs.jproteome.7b00572

Seok Ko, H., Uehara, T., Tsuruma, K., Nomura, Y., 2004. Ubiquilin interacts with ubiquitylated proteins and proteasome through its ubiquitin-associated and ubiquitin-like domains. FEBS Letters 566, 110–114. doi:10.1016/j.febslet.2004.04.031

Sharkey, L.M., Safren, N., Pithadia, A.S., Gerson, J.E., Dulchavsky, M., Fischer, S., Patel, R., Lantis, G., Ashraf, N., Kim, J.H., Meliki, A., Minakawa, E.N., Barmada, S.J., Ivanova, M.I., Paulson, H.L., 2018. Mutant UBQLN2 promotes toxicity by modulating intrinsic self-assembly. Proceedings of the National Academy of Sciences of the United States of America 115, E10495–E10504. doi:10.1073/pnas.1810522115

Shi, Y., Chen, X., Elsasser, S., Stocks, B.B., Tian, G., Lee, B.-H., Shi, Y., Zhang, N., de Poot, S.A.H., Tuebing, F., Sun, S., Vannoy, J., Tarasov, S.G., Engen, J.R., Finley, D., Walters, K.J., 2016. Rpn1 provides adjacent receptor sites for substrate binding and deubiquitination by the proteasome. Science 351. doi:10.1126/science.aad9421

Sims, J.J., Haririnia, A., Dickinson, B.C., Fushman, D., Cohen, R.E., 2009. Avid interactions underlie the Lys63-linked polyubiquitin binding specificities observed for UBA domains. Nat Struct Mol Biol 16, 883–889. doi:10.1038/nsmb.1637

Sreedharan, J., Blair, I.P., Tripathi, V.B., Hu, X., Vance, C., Rogelj, B., Ackerley, S., Durnall, J.C., Williams, K.L., Buratti, E., Baralle, F., de Belleroche, J., Mitchell, J.D., Leigh, P.N., Al-Chalabi, A., Miller, C.C., Nicholson, G., Shaw, C.E., 2008. TDP-43 Mutations in Familial and Sporadic Amyotrophic Lateral Sclerosis. Science 319, 1668–1672. doi:10.1126/science.1154584

Suzuki, R., Kawahara, H., 2016. UBQLN4 recognizes mislocalized transmembrane domain proteins and targets these to proteasomal degradation. EMBO Reports 17, 842–857. doi:10.15252/embr.201541402

Synofzik, M., Maetzler, W., Grehl, T., Prudlo, J., Hagen, Vom, J.M., Haack, T., Rebassoo, P., Munz, M., Schöls, L., Biskup, S., 2012. Screening in ALS and FTD patients reveals 3 novel UBQLN2 mutations outside the PXX domain and a pure FTD phenotype. Neurobiol. Aging 33, 2949.e13–7. doi:10.1016/j.neurobiolaging.2012.07.002

Şentürk, M., Lin, G., Zuo, Z., Mao, D., Watson, E., Mikos, A.G., Bellen, H.J., 2019. Ubiquilins regulate autophagic flux through mTOR signalling and lysosomal acidification. Nat. Cell Biol. 21, 384–396. doi:10.1038/s41556-019-0281-x

Teyssou, E., Chartier, L., Amador, M.-D.-M., Lam, R., Lautrette, G., Nicol, M., Machat, S., Da Barroca, S., Moigneu, C., Mairey, M., Larmonier, T., Saker, S., Dussert, C., Forlani, S., Fontaine, B., Seilhean, D., Bohl, D., Boillée, S., Meininger, V., Couratier, P., Salachas, F., Stevanin, G., Millecamps, S., 2017. Novel UBQLN2 mutations linked to amyotrophic lateral sclerosis and atypical hereditary spastic paraplegia phenotype through defective HSP70-mediated proteolysis. Neurobiol. Aging 58, 239.e11– 239.e20. doi:10.1016/j.neurobiolaging.2017.06.018

Tsuchiya, H., Ohtake, F., Arai, N., Kaiho, A., Yasuda, S., Tanaka, K., Saeki, Y., 2017. In Vivo Ubiquitin Linkage-type Analysis Reveals that the Cdc48-Rad23/Dsk2 Axis Contributes to K48-Linked Chain Specificity of the Proteasome. Molecular Cell 66, 488–502.e7. doi:10.1016/j.molcel.2017.04.024

Vermeiren, Y., Janssens, J., Van Dam, D., De Deyn, P.P., 2018. Serotonergic Dysfunction in Amyotrophic Lateral Sclerosis and Parkinson’s Disease: Similar Mechanisms, Dissimilar Outcomes. Front Neurosci 12, 185. doi:10.3389/fnins.2018.00185

Walters, K.J., Kleijnen, M.F., Goh, A.M., Wagner, A., Howley, P.M. 2002. Structural Studies of the Interaction between Ubiquitin Family Proteins and Proteasome Subunit S5a†. Biochemistry 41, 1767–1777. doi:10.1021/bi011892y

Wang, H., Lim, P.J., Yin, C., Rieckher, M., Vogel, B.E., Monteiro, M.J., 2006. Suppression of polyglutamine-induced toxicity in cell and animal models of Huntington’s disease by ubiquilin. Human Molecular Genetics 15, 1025–1041. doi:10.1093/hmg/ddl017

Weber, M., Wu, T., Hanson, J.E., Alam, N.M., Solanoy, H., Ngu, H., Lauffer, B.E., Lin, H.H., Dominguez, S.L., Reeder, J., Tom, J., Steiner, P., Foreman, O., Prusky, G.T., Scearce-Levie, K., 2015. Cognitive Deficits, Changes in Synaptic Function, and Brain Pathology in a Mouse Model of Normal Aging(1,2,3). eNeuro 2, ENEURO.0047– 15.2015. doi:10.1523/ENEURO.0047-15.2015

Whiteley, A.M., Prado, M.A., Peng, I., Abbas, A.R., Haley, B., Paulo, J.A., Reichelt, M., Katakam, A., Sagolla, M., Modrusan, Z., Lee, D.Y., Roose-Girma, M., Kirkpatrick, D.S., McKenzie, B.S., Gygi, S.P., Finley, D., Brown, E.J., 2017. Ubiquilin1 promotes antigen-receptor mediated proliferation by eliminating mislocalized mitochondrial proteins. eLife Sciences 6, e26435. doi:10.7554/eLife.26435

Wilkinson, C.R., Seeger, M., Hartmann-Petersen, R., Stone, M., Wallace, M., Semple, C., Gordon, C., 2001. Proteins containing the UBA domain are able to bind to multi-ubiquitin chains. Nat. Cell Biol. 3, 939–943. doi:10.1038/ncb1001-939

Williams, K.L., Warraich, S.T., Yang, S., Solski, J.A., Fernando, R., Rouleau, G.A., Nicholson, G.A., Blair, I.P., 2012. UBQLN2/ubiquilin 2 mutation and pathology in familial amyotrophic lateral sclerosis. Neurobiol. Aging 33, 2527.e3–10. doi:10.1016/j.neurobiolaging.2012.05.008

Wood, J.D., Beaujeux, T.P., Shaw, P.J., 2003. Protein aggregation in motor neurone disorders. Neuropathology and Applied Neurobiology 29, 529–545. doi:10.1046/j.0305-1846.2003.00518.x

Yokota, T., Mishra, M., Akatsu, H., Tani, Y., Miyauchi, T., Yamamoto, T., Kosaka, K., Nagai, Y., Sawada, T., Heese, K., 2006. Brain site-specific gene expression analysis in Alzheimer’s disease patients. European Journal of Clinical Investigation 36, 820– 830. doi:10.1111/j.1365-2362.2006.01722.x

Yu, G., Wang, L.-G., Han, Y., He, Q.-Y., 2012. clusterProfiler: an R package for comparing biological themes among gene clusters. OMICS 16, 284–287. doi:10.1089/omi.2011.0118

Zhang, D., Raasi, S., Fushman, D., 2008. Affinity Makes the Difference: Nonselective Interaction of the UBA Domain of Ubiquilin-1 with Monomeric Ubiquitin and Polyubiquitin Chains. Journal of Molecular Biology 377, 162–180. doi:10.1016/j.jmb.2007.12.029

Zhang, Y., Pak, C., Han, Y., Ahlenius, H., Zhang, Z., Chanda, S., Marro, S., Patzke, C., Acuna, C., Covy, J., Xu, W., Yang, N., Danko, T., Chen, L., Wernig, M., Südhof, T.C., 2013. Rapid single-step induction of functional neurons from human pluripotent stem cells. Neuron 78, 785–798. doi:10.1016/j.neuron.2013.05.029

Zuris, J.A., Thompson, D.B., Shu, Y., Guilinger, J.P., Bessen, J.L., Hu, J.H., Maeder, M.L., Joung, J.K., Chen, Z.-Y., Liu, D.R., 2015. Cationic lipid-mediated delivery of proteins enables efficient protein-based genome editing in vitro and in vivo. Nat. Biotechnol. 33, 73–80. doi:10.1038/nbt.3081

