## Supplemental Figures for "Global proteomics of *Ubqln*2-based murine models of ALS"

Supplemental Figure 1

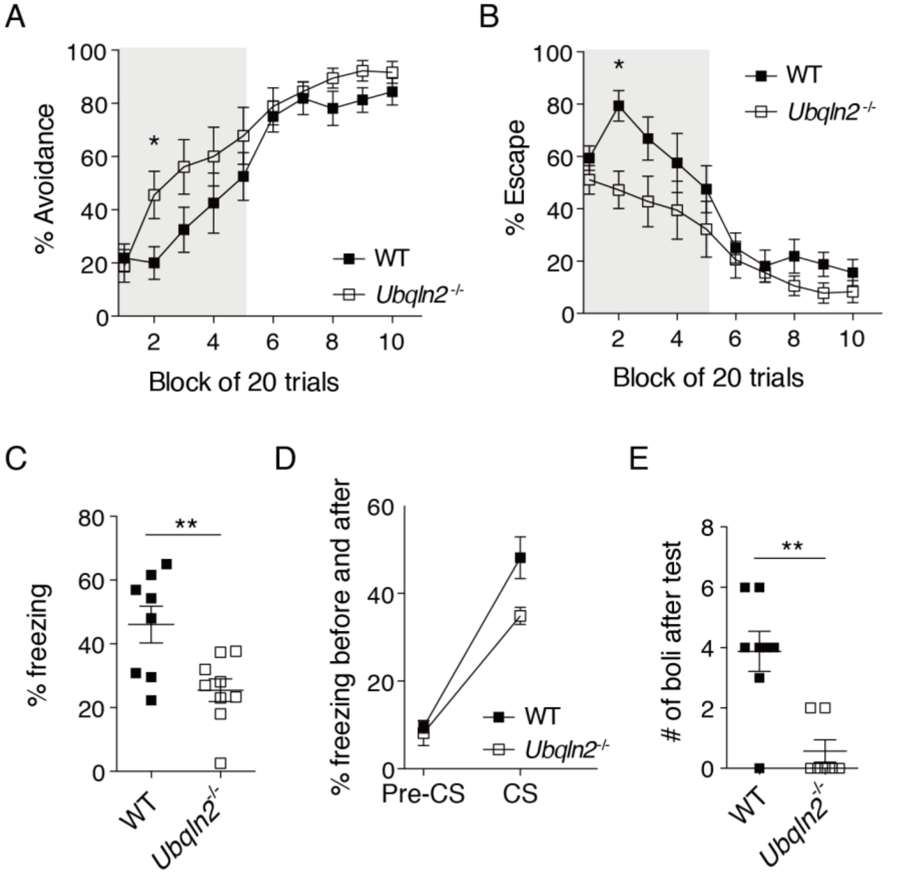


**Supplemental Figure 1: *Ubqln2-/-* mice have memory deficits.** (A, B) *Ubqln2-/-* mice exhibit largely normal active avoidance behavior. (C) *Ubqln2-/-* mice exhibit reduced % freezing during the contextual fear conditioning test. (D) *Ubqln2-/-* mice have a mild decrease in % freezing during the presentation of the CS in the cued fear conditioning test. (E) The number of fecal boli left in the chamber after the cued-fear condition test period was used as a proxy measure of anxiety and emotionality. *Ubqln2-/-* mice had less boli when compared to WT controls. Statistical significance was determined with two-tailed, unpaired Student's T test. N=8-9 animals.

Supplemental Figure 2

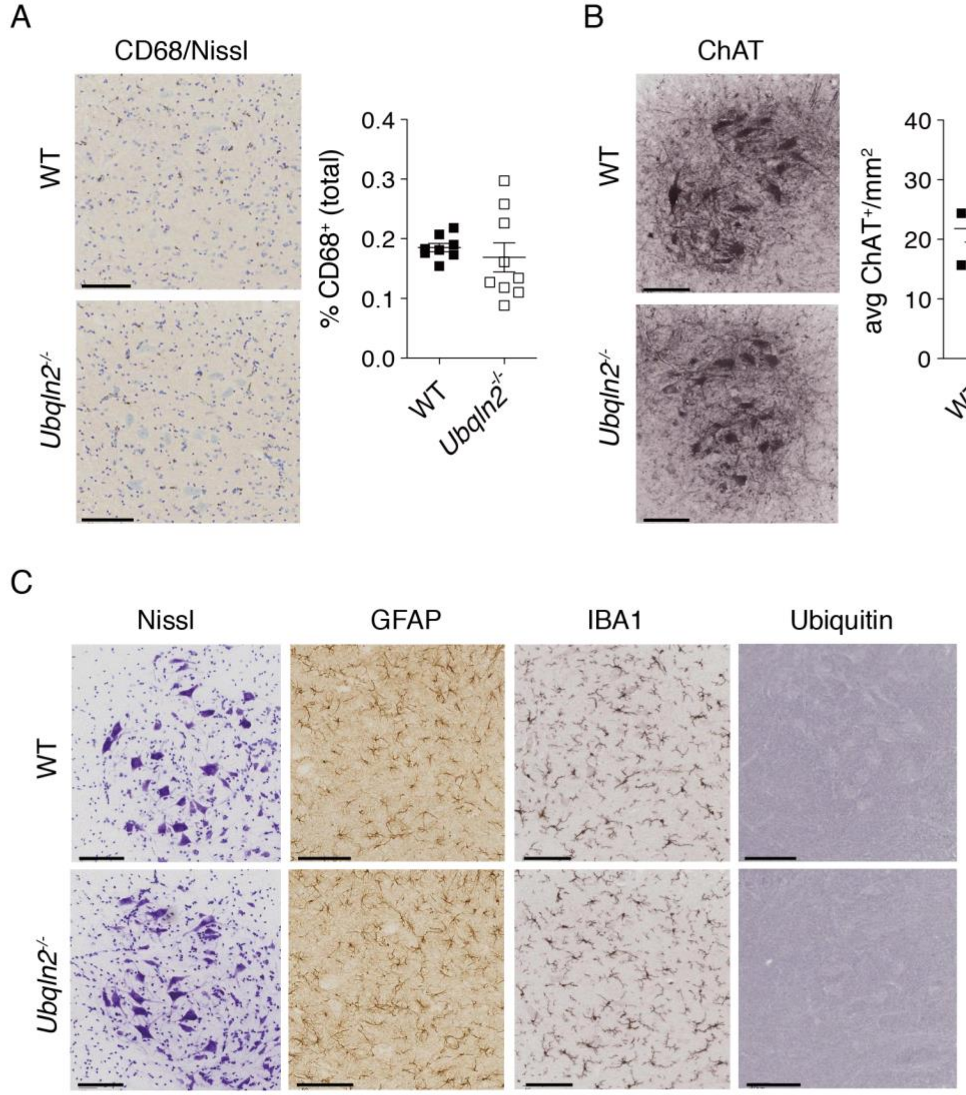


**Supplemental Figure 2: Histological staining of *Ubqln2-/-* animals.** Spinal cords of approximately one-year old mice were harvested for histological staining and quantitation. (A) CD68 and Nissl co-staining reveals no changes in neuroinflammation. (B) *Ubqln2-/-* spinal cords have normal ChAT+ neuron quantitation. (C) No significant changes in Nissl, GFAP, IBA1, or Ubiquitin staining. N=8-9 animals per genotype. Scale bar = 100 μm for each image.

Supplemental Figure 3

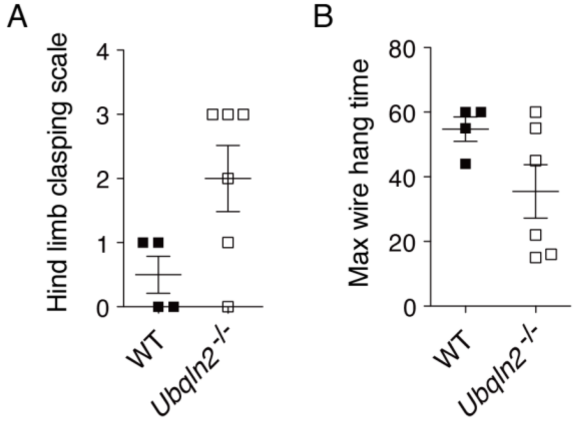


**Supplemental Figure 3: Neuromotor testing of young animals.** Approximately 4-5 month-old animals were tested for hind limb clasping (A) and maximum wire hang time (B) as in Figure 1. N=4-6 animals per genotype, and statistical significance was determined via unpaired, two-tailed Student's T test.

Supplemental Figure 4

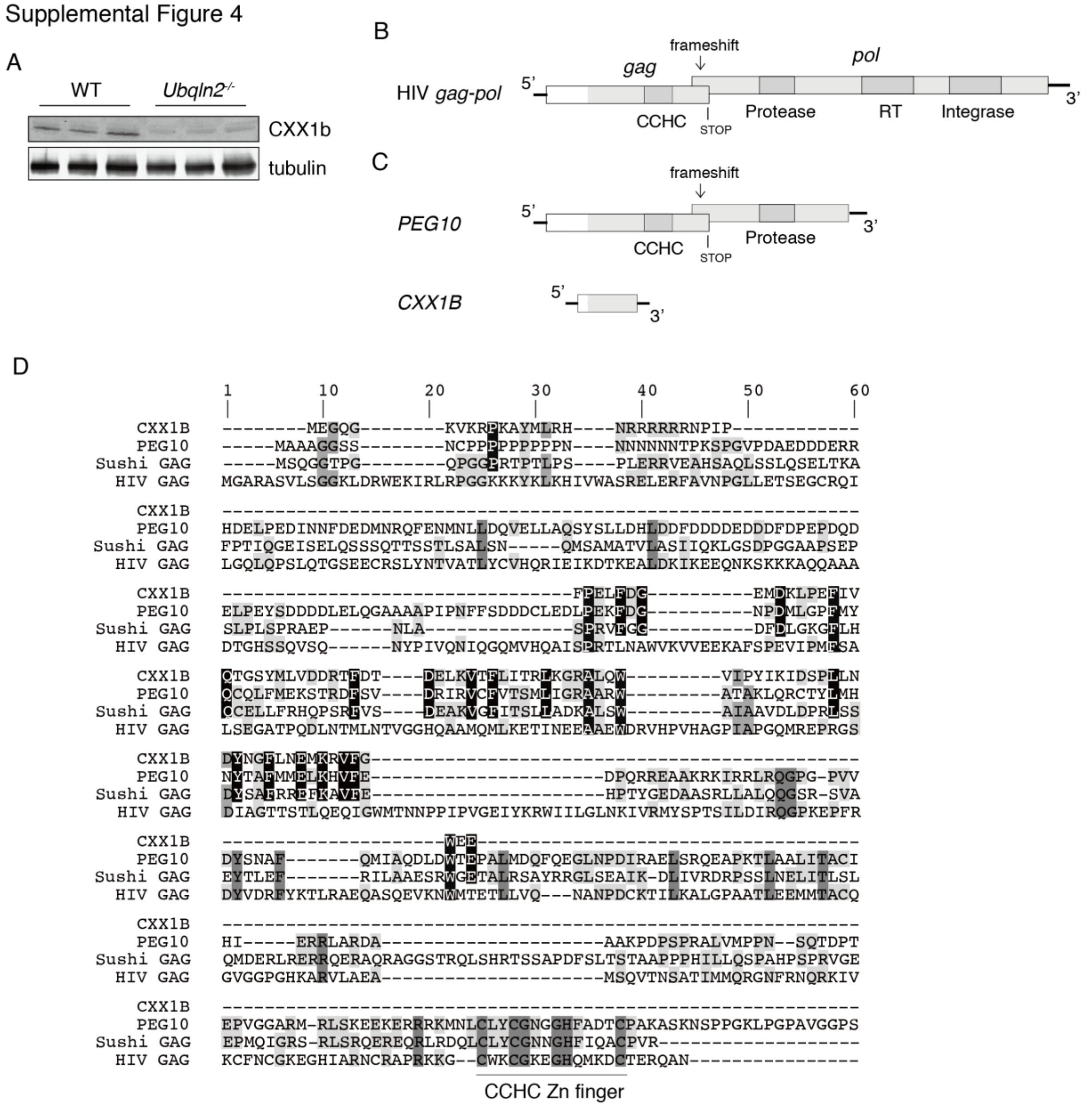


**Supplemental Figure 4: Further focus on initial proteomics hits from *Ubqln2-/-* animals.** (A) Spinal cord of 6-month old WT and *Ubqln2-/-* animals was lysed in 8 M urea buffer and prepared for Western blot of CXX1B. (B) Schematic of *gag*-*pol*-containing retroelement genes. Early stop codon is denoted with 'STOP', which is in close proximity to a slippery mRNA sequence that results in frequent frameshifting and translation of a second reading frame contiguous with the first. Functional domains of the *gag* and *pol* regions (in light grey) are boxed in dark grey. (C) Detailed mRNA schematic of the Ty3/Gypsy retroelement family members *Peg10* and *Cxx1b* from Figure 2. Top: *Peg10* mRNA organization showing functional protein domains in dark grey boxes. Bottom: *Cxx1b* mRNA organization with annotated *gag* domain. (D) Amino acid alignment of CXX1B, murine GAG domain of PEG10, ancestral Sushi-ichi GAG (AAC33525), and HIV

GAG (AAA76686). Amino acids with 100% conservation amongst Ty3/gypsy retroelement family members are highlighted in black. Amino acids conserved amongst PEG10, Sushi GAG, and HIV GAG in the absence of CXX1B alignment are highlighted in dark grey; amino acids conserved amongst three of four aligned sequences in medium grey, and amino acids conserved among only two aligned sequences in light grey. CCHC domain of PEG10 and Sushi/HIV GAG proteins is noted at the bottom. Alignment was performed using MUSCLE algorithm in Geneious with 8 iterations.

Supplemental Figure 5

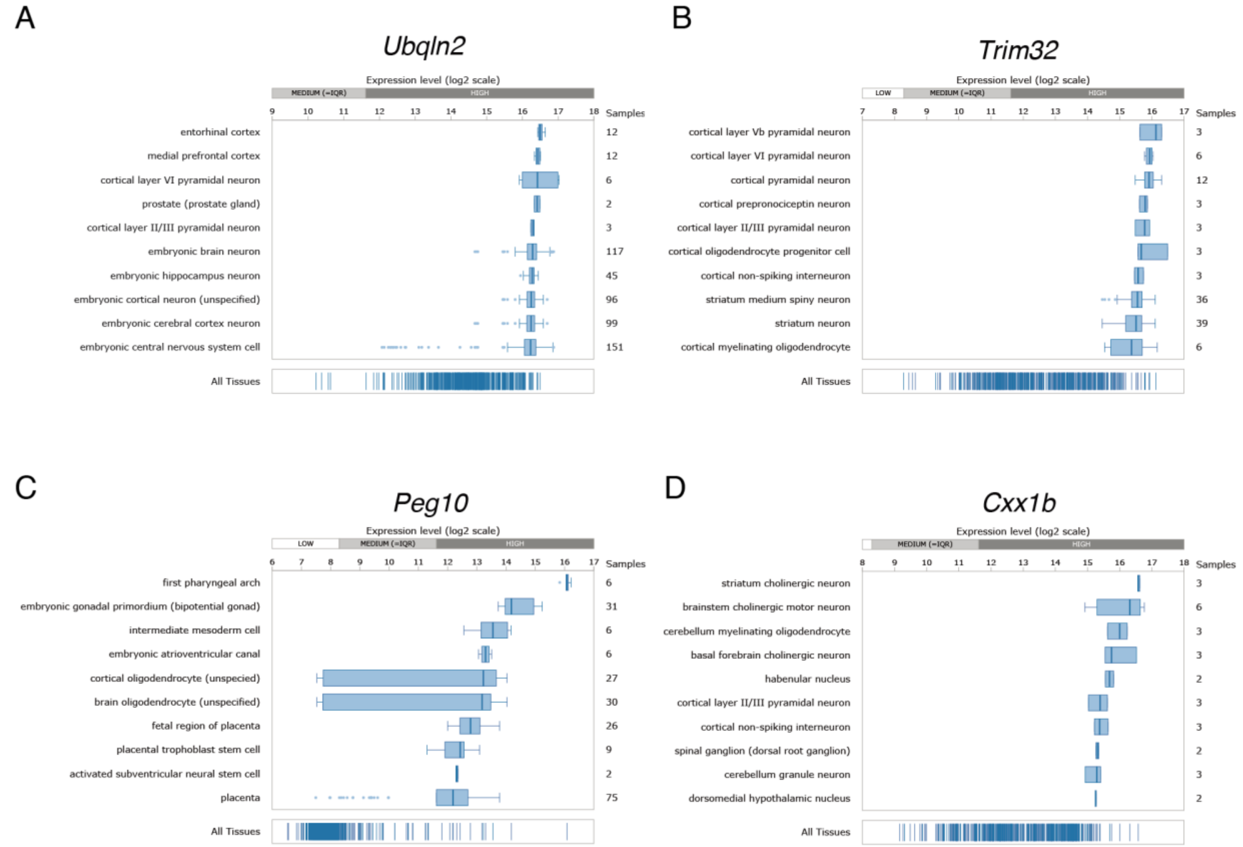


**Supplemental Figure 5: Gene expression in multiple mouse tissues.** (A) *Ubqln2*, (B) *Trim32*, (C) *Peg10* and (D) *Cxx1b* mRNA are expressed at relatively high levels in neuronal tissues. Data are extracted from Genevestigator (Hruz et al., 2008), a search engine that allows the comparison and analysis of the transcriptional regulation of single genes across multiple species, tissues and conditions.

Supplemental Figure 6

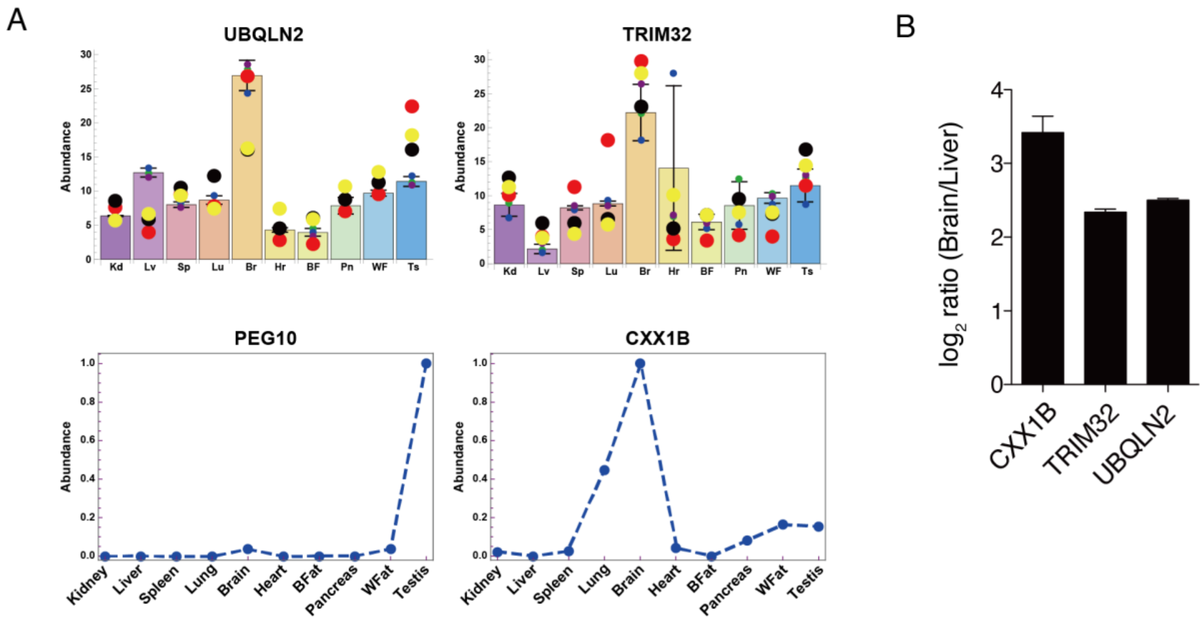


**Supplemental Figure 6: Protein expression in multiple mouse tissues.** (A) Protein expression of hits in 10 different tissues of mice. TRIM32, CXX1B and UBQLN2 are most abundant in brain. The two upper graphs represent TMT-MS3 data. Lower graphs are label free quantifications. Kd, Kidney; Lv, Liver; Sp, Spleen; Lu, Lung; Br, Brain; Hr, Heart; BF, Brown Fat; Pn, Pancreas; WF, White fat; Ts, Testis. (B) Comparison of brain and liver proteomes; CXX1B, TRIM32 and UBQLN2 are more abundant in the brain. PEG10 was not quantified in this dataset. Data extracted from (Rose et al., 2016).

Supplemental Figure 7

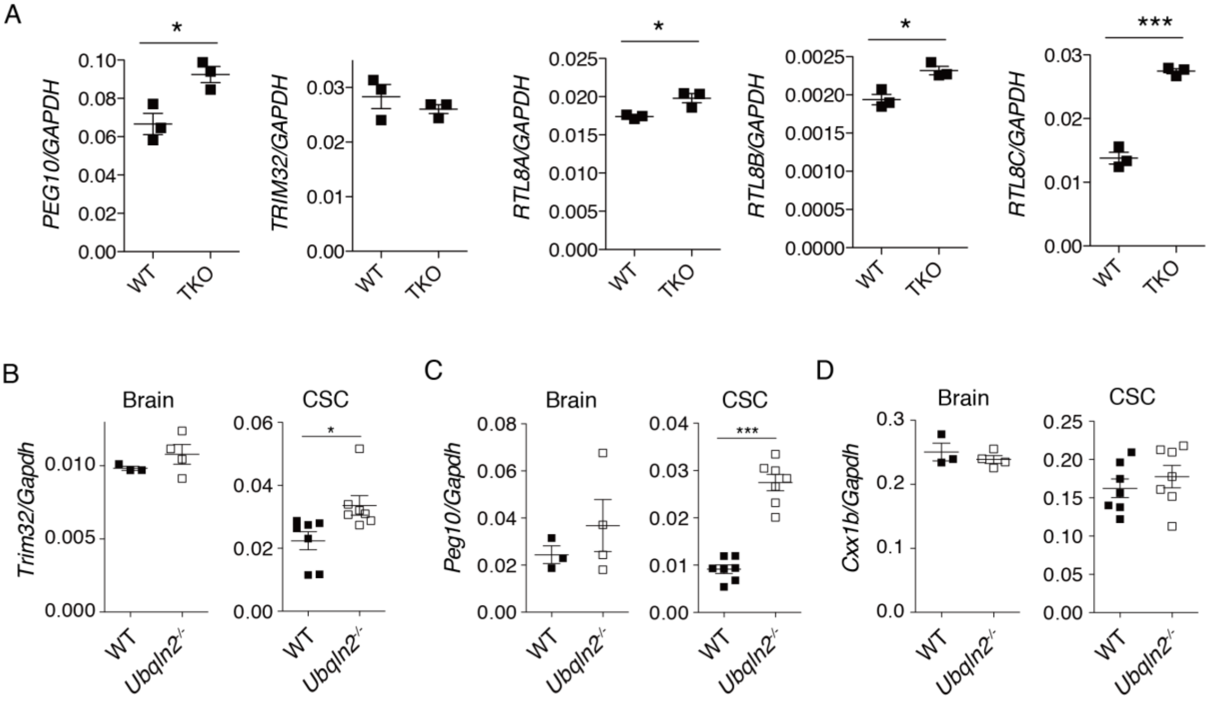


**Supplemental Figure 7: mRNA levels of top hits in HEK 293 cells and in mouse tissues.** mRNA abundance of shared proteomic hits from Figures 2 and 3. (A) mRNA of endogenous human *PEG10*, *TRIM32*, and *RTL8A-C* in WT and TKO HEK 293 cells. N=3 independent experiments. (B) mRNA of murine *Trim32* (B), *Peg10* (C), and *Cxx1b* (D) as measured by QPCR of brain (left, at approximately 1 year of age) and cervical spinal cord (CSC, right, at approximately 5-6 months of age) of WT and *Ubqln2*$^{-/-}$ animals. Statistics were determined via unpaired, two-tailed Student's T test. N=3-4 animals per genotype for brain and 7 animals per genotype for spinal cord.

Supplemental Figure 8

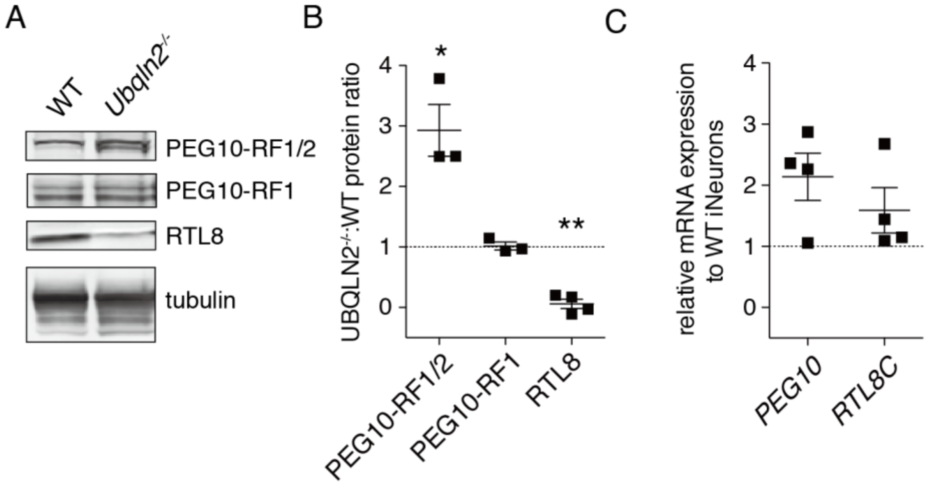


**Supplemental Figure 8: Quantifying alteration of top hits in human iNeurons.** (A) Example Western blot of PEG10-RF1/2, PEG10-RF1, and CXX1B in human induced neurons (iNeurons) after 7 days of differentiation from ES cell precursors. (B) Quantification of the protein abundance of the top hits in *Ubqln2-/-* iNeurons compared to WT controls (normalized to 1 for each unique differentiation experiment). Protein abundance is first normalized to tubulin levels, then *Ubqln2-/-* iNeuron protein levels are normalized to WT expression levels of the same proteins. (C) Quantification of mRNA levels of top hits from (A) as measured by QPCR with gene expression normalized to GAPDH. Following GAPDH normalization, *Ubqln2-/-* iNeuron mRNA levels are normalized to WT expression levels of the same genes. Statistical significance was determined via a paired, two-tailed Student’s T test.

Supplemental Figure 9

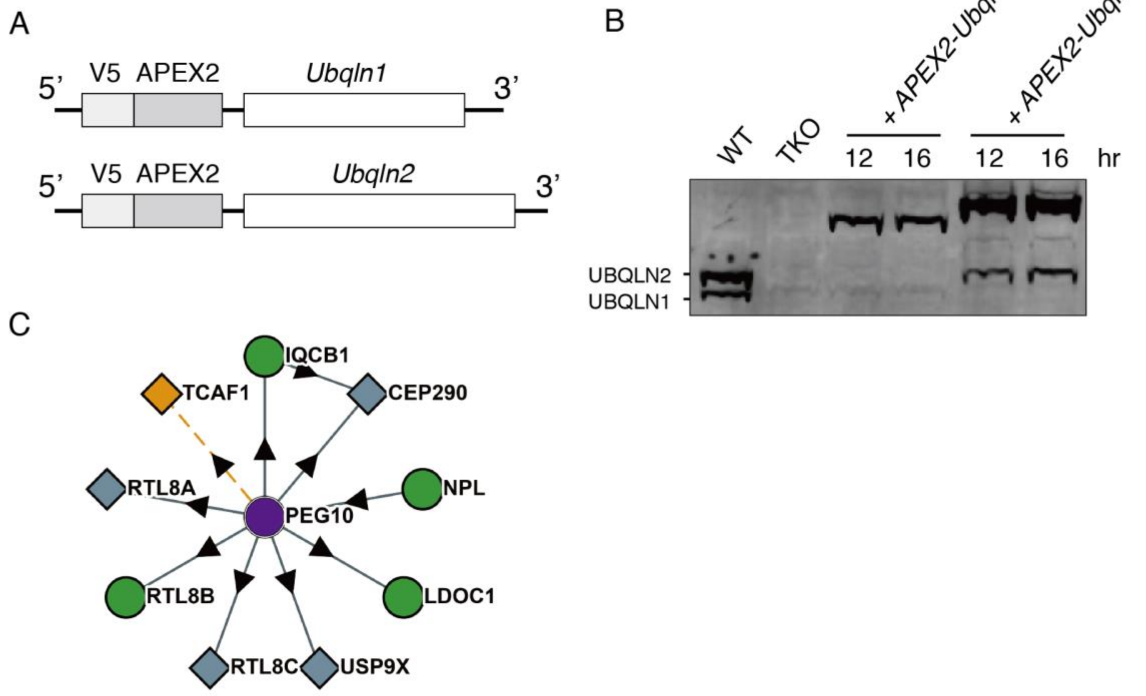


**Supplemental Figure 9: Setup of APEX2 proximity-labelling experiment.** (A) Schematic of APEX2-Ubqln fusion constructs used in experiment. (B) Western blot of endogenous UBQLN1 and UBQLN2 in WT HEK 293 cells, compared to loss of endogenous protein in TKO HEK 293 cells and expression of APEX2-fusion proteins. (C) Interactome analysis of human PEG10, adapted from BioPlex (Huttlin et al., 2017). Purple circle is queried protein (PEG10); green circles are bait proteins; gray diamonds are prey proteins; orange square represents subthreshold interactor. Directed edge (indicated by an arrow) follows bait-to-prey testing relationship.

Supplemental Figure 10

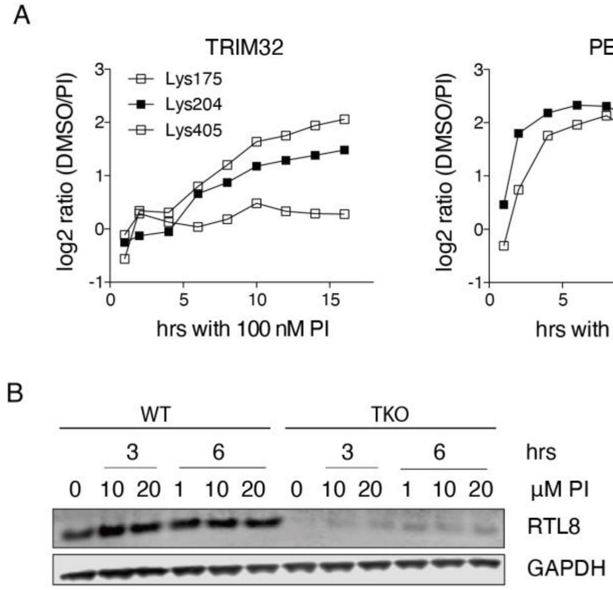


**Supplemental Figure 10: Quantifying accumulation of top hits upon inhibition of protein degradation.** (A) GlyGly sites quantified for TRIM32 (left), PEG10 (middle), and RTL8/FAM127B (right) after 100 nM bortezomib and epoxomycin (PI) treatment. Different symbols reflect unique ubiquitinated peptides quantified by GyGly-TMT analysis. Data are adapted from (Rose et al., 2016). (B) WT or TKO HEK cells were incubated with high doses of proteasome inhibitors (PI, 1 to 20 µM) for either three or six hr and probed for RTL8 protein levels. GAPDH was used as loading control.

Supplemental Figure 11

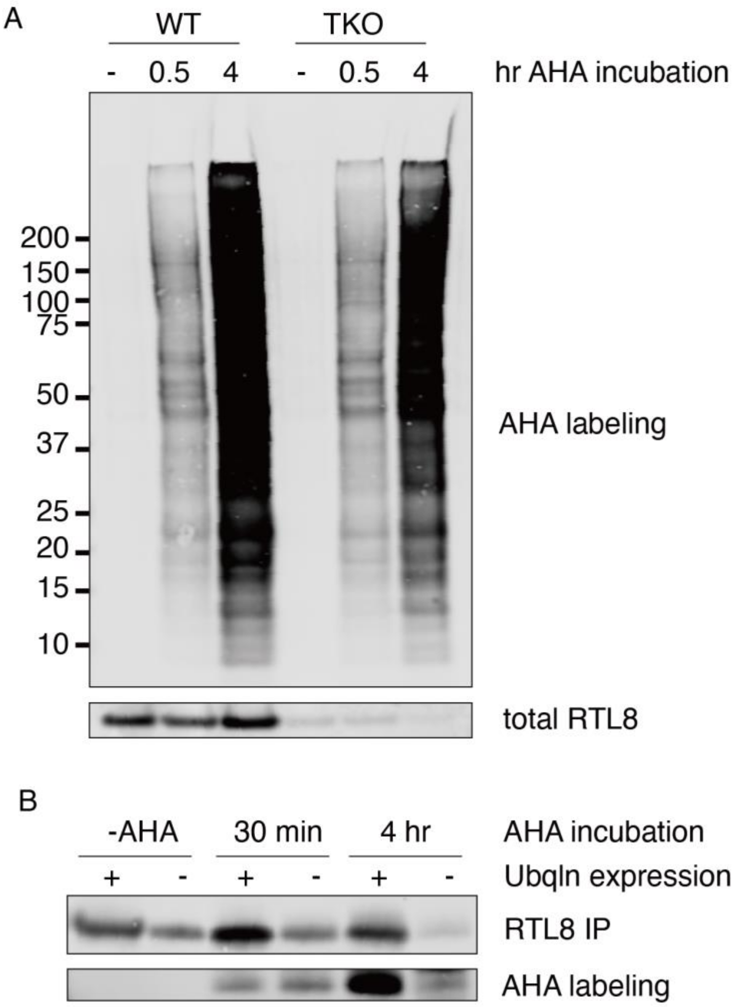


**Supplemental Figure 11: RTL8 synthesis in Ubqln-deficient cells.** WT or TKO HEK 293 cells were incubated with the methionine analog AHA in order to label nascent protein. Following AHA incubation, RTL8 was immunoprecipitated to quantify protein synthesis. (A) AHA labeling of whole cell lysate. Cells were incubated with regular (-) or AHA-containing media for 30 min or 4 hr following 45 min of methionine starvation. Following lysis, nascent protein was labelled with biotin via click-chemistry and probed with streptavidin. Endogenous total RTL8 levels were also quantified. (B) AHA labeling of immunoprecipitated RTL8 protein. RTL8 was immunoprecipitated from AHA-labelled lysate of $10_7$ cells per condition, labelled with biotin via click-chemistry, and the proportion of total RTL8 was determined via streptavidin probe. Shown is one of two representative experiments.

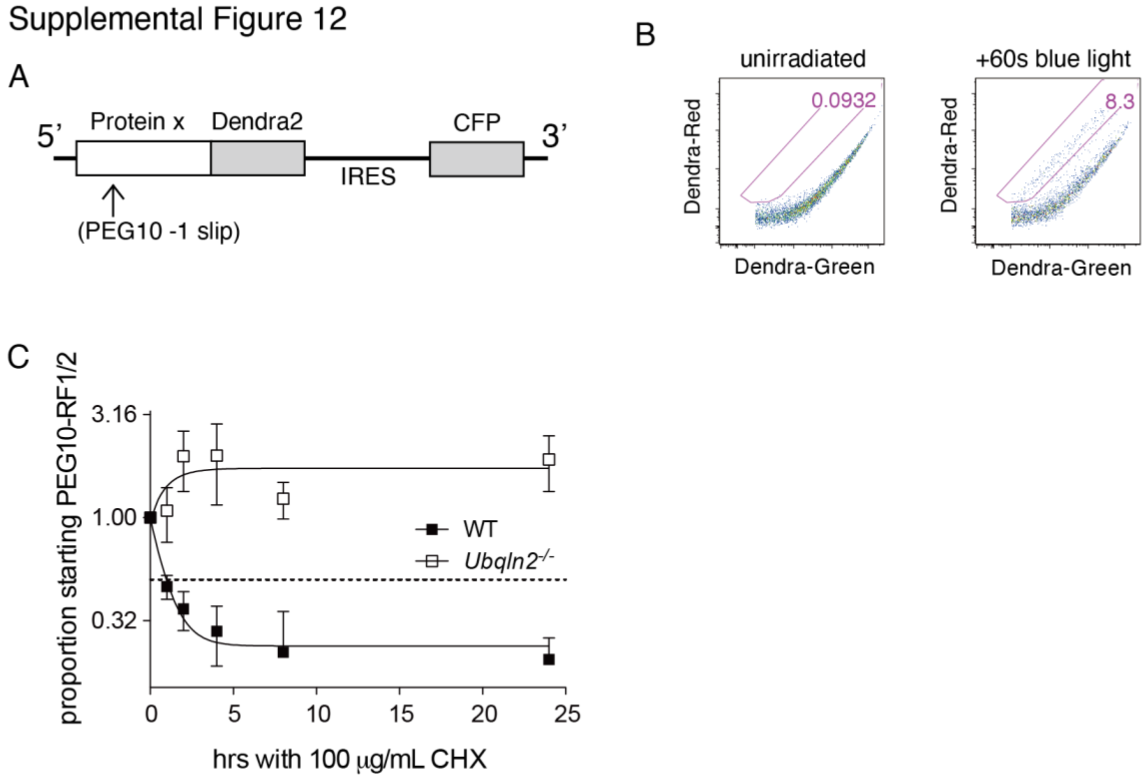


**Supplemental Figure 12: Setup of Dendra2 time course experiment.** (A) Construct design of putative UBQLN2 client proteins. *Peg10*'s premature stop codon is highlighted with arrow indicating the -1 slip site before the insertion of a Dendra2 cassette. (B) Representative flow cytometry dot plots demonstrating GFP-Dendra2 signal and photoconversion in transfected cells. Green Dendra2 fluorescence was observed with Alexa Flour 488 filter, and red Dendra2 fluorescence was observed with PE filter. Cells that become positive for red Dendra2 signal were gated and the percentage of cells with photoconverted protein is shown in pink. (C) Degradation rate of endogenous full-length PEG10 protein in human iNeurons. Cells were treated with 100 μg/mL cycloheximide and samples were taken at the indicated timepoints for Western blot. N=4 independent experiments.

Supplemental Figure 13

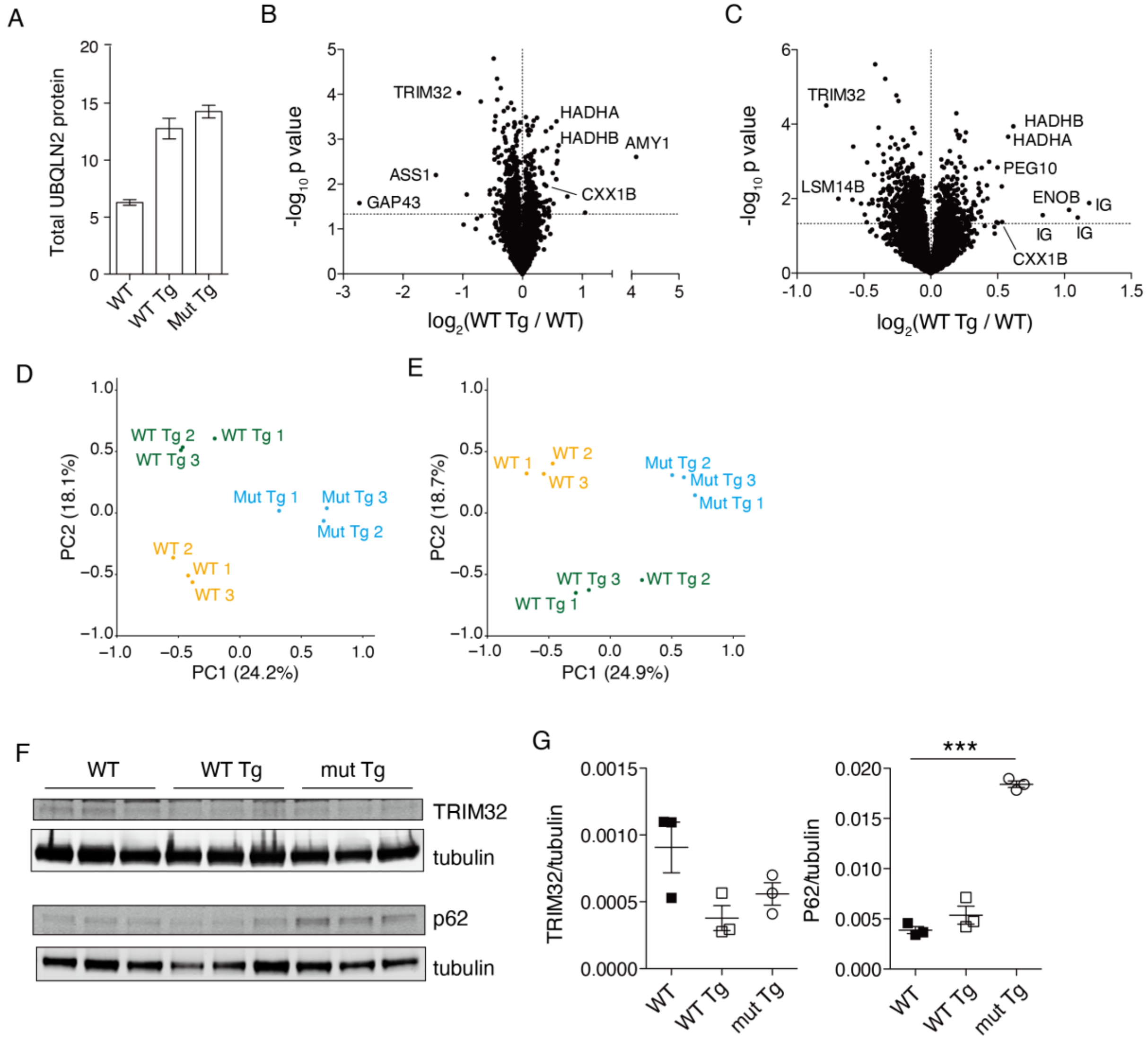


**Supplemental Figure 13: Proteomic results from transgenic *Ubqln2* overexpression mice.** (A) Quantification of total combined mouse and human UBQLN2 in transgenic mice. Two peptides shared amongst mouse and human UBQLN2 were quantified by TMT 10-plex in hippocampal tissue between WT, WT Tg, and Mutant Tg mice. (B-C) Hippocampus (B) and lumbar spinal cord (C) protein from transgenic mice expressing WT *Ubqln2* (WT Tg) was compared to WT littermates lacking transgene expression (WT). N=3 animals per genotype. (D-E) PCA analysis of transgenic hippocampus (D) and spinal cord (E) samples. Transgene-negative (WT) mice are in yellow, WT transgene-overexpressing mice are in green, and mutant transgene-overexpressing mice are in blue. (F) Protein samples from *Ubqln2* transgenic mice were prepared as in Figure 5 and Supplemental Figure 4 for Western blot confirmation of protein changes, including

TRIM32 and SQSTM1/P62. (G) Quantitation of (F) following normalization to tubulin. Statistical significance was determined by using an unpaired, two-tailed Student’s T test.
