## Supplemental Table 2 for "Global proteomics of *Ubqln*2-based murine models of ALS"

| **TMT** | **Experiment** | **Ubqln2** | **Tissue** | **Age (months)** | **Sex** | **Run ID** | **MS Instrument** |
| --- | --- | --- | --- | --- | --- | --- | --- |
| **1** | KO Old Brain | KO | Brain | 12-16 | Male | m07953-62 | Obitrap Fusion |
| **2** | KO Young Brain | KO | Brain | 5-6 | Male | a01805-16 | Obitrap Lumos |
| **3** | KO Lumb SpC | KO | Lumbar Spinal Cord | 5-6 | Male | a01833-44 | Obitrap Lumos |
| **4** | KO Hippo | KO | Hippocampus | 4-4.5 | Male | m11795-806 | Obitrap Fusion |
| **5** | KI Hippo | KI | Hippocampus | 4-5 | Male | m09995-10006 | Obitrap Fusion |
| **6** | KI SpC | KI | Spinal Cord | 4-5 | Male | m10008-19 | Obitrap Fusion |
| **7** | Tg Hippo | WT-hTg/Mut-hTg | Hippocampus | 2 | 8M 1F | m14899-10 | Obitrap Fusion |
| **8** | Tg Lumb SpC | WT-hTg/Mut-hTg | Lumbar Spinal Cord | 2 | 8M 1F | m14925-36 | Obitrap Fusion |

**TMT_1: *Ubqln2* KO OLD Brain:**

| **TMT label** | **Animal ID** | **Sample** | **Sex** | **Neuromotor testing age (days)** | **Neuromotor testing weight (g)** | **Tissue harvest age (days)** | ***Ubqln2*** |
| --- | --- | --- | --- | --- | --- | --- | --- |
| 126 | 5114-461 | WT_1 | M | 314 | 31.5 | 365 | wt/y |
| 127N | 5114-460 | WT_2 | M | 314 | 37 | 365 | wt/y |
| 127C | 5114-463 | WT_3 | M | 314 | 37.4 | 365 | wt/y |
| 128N | 5114-394 | WT_4 | M | 448 | 33.4 | 497 | wt/y |
| 128C | 5114-395 | WT_5 | M | 448 | 32.4 | 497 | wt/y |
| 129N | 5114-449 | KO_1 | M | 321 | 35.1 | 370 | ko/y |
| 129C | 5114-445 | KO_2 | M | 321 | 37.5 | 370 | ko/y |
| 130N | 5114-446 | KO_3 | M | 321 | 32.6 | 370 | ko/y |
| 130C | 5114-385 | KO_4 | M | 448 | 31.3 | 497 | ko/y |
| 131 | 5114-388 | KO_5 | M | 448 | 31 | 497 | ko/y |

**TMT_2_and_3: *Ubqln2* KO Young (Brain and Lumbar Spinal Cord):**

| **TMT label** | **Animal ID** | **Sample** | **Sex** | **Neuromotor testing age (days)** | **Neuromotor testing weight (g)** | **Tissue harvest age (days)** | ***Ubqln2*** |
| --- | --- | --- | --- | --- | --- | --- | --- |
| 126 | 5114-728 | WT_1 | M | 137 | 28 | 143 | wt/y |
| 127N | 5114-738 | WT_2 | M | 137 | 30.5 | 143 | wt/y |
| 127C | 5114-741 | WT_3 | M | 137 | 29.8 | 143 | wt/y |
| 128N | 5114-743 | WT_4 | M | 137 | 31.3 | 143 | wt/y |
| 128C | 5114-712 | KO_1 | M | 170 | 26.6 | 176 | ko/y |
| 129N | 5114-718 | KO_2 | M | 165 | 26 | 171 | ko/y |
| 129C | 5114-731 | KO_3 | M | 137 | 31.5 | 143 | ko/y |
| 130N | 5114-732 | KO_4 | M | 137 | 27.9 | 143 | ko/y |
| 130C | 5114-737 | KO_5 | M | 137 | 30.1 | 143 | ko/y |
| 131 | 5114-742 | KO_6 | M | 137 | 31.8 | 143 | ko/y |

**TMT_4: *Ubqln2* KO Hippocampus:**

| **TMT Label** | **Animal ID** | **Sample** | **Sex** | **Age (days)** | ***Ubqln2*** | **Hippocampus** |
| --- | --- | --- | --- | --- | --- | --- |
| 126 | 5114-919 | WT_1 | M | 106 | wt/y | Right |
| 127N | 5114-918 | WT_2 | M | 106 | wt/y | Right |
| 127C | 5114-877 | WT_3 | M | 131 | wt/y | Right |
| 128N | 5114-922 | KO_1 | M | 106 | ko/y | Right |
| 128C | 5114-920 | KO_2 | M | 106 | ko/y | Right |
| 129N | 5114-919 | WT_1 | M | 106 | wt/y | Left |
| 129C | 5114-918 | WT_2 | M | 106 | wt/y | Left |
| 130N | 5114-922 | KO_1 | M | 106 | ko/y | Left |
| 130C | 5114-920 | KO_2 | M | 106 | ko/y | Left |
| 131 | 5114-878 | KO_3 | M | 131 | ko/y | Left |

**TMT_5: *Ubqln2* KI Hippocampus:**

| **TMT Label** | **Animal ID** | **Sample** | **Sex** | **Age (days)** | ***Ubqln2*** | **Hippocampus** |
| --- | --- | --- | --- | --- | --- | --- |
| 126 | 289067 | WT_1 | M | 132 | wt | Left |
| 127N | 292049 | WT_2 | M | 112 | wt | Left |
| 127C | 289790 | WT_3 | M | 121 | wt | Left |
| 128N | 290262 | KI_1 | M | 114 | P520T | Left |
| 128C | 290260 | KI_2 | M | 114 | P520T | Left |
| 129N | 289067 | WT_4 | M | 132 | wt | Right |
| 129C | 292049 | WT_5 | M | 112 | wt | Right |
| 130N | 289790 | WT_6 | M | 121 | wt | Right |
| 130C | 290262 | KI_3 | M | 114 | P520T | Right |
| 131 | 290260 | KI_4 | M | 114 | P520T | Right |

**TMT_6: *Ubqln2* KI Spinal Cord:**

| **TMT Label** | **Animal ID** | **Sample** | **Sex** | **Age (days)** | ***Ubqln2*** | **Spinal Cord** |
| --- | --- | --- | --- | --- | --- | --- |
| 126 | 289067 | WT_1 | M | 132 | wt | Lumbar |
| 127N | 292049 | WT_2 | M | 112 | wt | Lumbar |
| 127C | 289790 | WT_3 | M | 121 | wt | Lumbar |
| 128N | 290262 | KI_1 | M | 114 | P520T | Lumbar |
| 128C | 290260 | KI_2 | M | 114 | P520T | Lumbar |
| 129N | 289067 | WT_4 | M | 132 | wt | Cervical |
| 129C | 292049 | WT_5 | M | 112 | wt | Cervical |
| 130N | 289790 | WT_6 | M | 121 | wt | Cervical |
| 130C | 290262 | KI_3 | M | 114 | P520T | Cervical |
| 131 | 290260 | KI_4 | M | 114 | P520T | Cervical |

**TMT_7_and_8: *Ubqln2* hTg Hippocampus and Lumbar Spinal Cord:**

| **TMT label** | **Animal ID** | **Sample** | **Sex** | **Age (days)** | ***Ubqln2*** |
| --- | --- | --- | --- | --- | --- |
| 126 | 1194 | WT_Tg_1 | M | 60 | WT_hTg |
| 127N | 1197 | WT_Tg_2 | M | 60 | WT_hTg |
| 127C | 1191 | WT_Tg_3 | F | 60 | WT_hTg |
| 128N | 1178 | Mut_Tg_1 | M | 63 | P497S_hTg |
| 128C | 1182 | Mut_Tg_2 | M | 63 | P497S_hTg |
| 129N | 1183 | Mut_Tg_3 | M | 63 | P497S_hTg |
| 129C | 1179 | No_Tg_1 | M | 63 | noTg |
| 130N | 1180 | No_Tg_2 | M | 63 | noTg |
| 130C | 1181 | No_Tg_3 | M | 63 | noTg |
